# Cell environment shapes TDP-43 function: implications in neuronal and muscle disease

**DOI:** 10.1101/2021.04.20.440589

**Authors:** Urša Šušnjar, Neva Škrabar, Anna-Leigh Brown, Yasmine Abbassi, NYGC ALS Consortium, Hemali Phatnani, Andrea Cortese, Cristina Cereda, Enrico Bugiardini, Rosanna Cardani, Giovanni Meola, Michela Ripolone, Maurizio Moggio, Maurizio Romano, Maria Secrier, Pietro Fratta, Emanuele Buratti

**Affiliations:** Molecular Pathology Lab, International Centre for Genetic Engineering and Biotechnology (ICGEB), Trieste, Italy; Tumour Virology Lab, International Centre for Genetic Engineering and Biotechnology (ICGEB), Trieste, Italy; Generatio GmbH, Center for Animal, Genetics, Tübingen, Germany; Department of Neuromuscular Diseases, UCL Queen Square Institute of Neurology, London, UK; Center for Genomics of Neurodegenerative Disease, New York Genome Center, New York, USA; Department of Brain and Behaviour Sciences, University of Pavia, Pavia, Italy; Genomic and post-Genomic Unit, IRCCS Mondino Foundation, Pavia, Italy; BioCor Biobank, UOC SMEL-1 of Clinical Pathology, IRCCS-Policlinico San Donato, San Donato Milanese, Italy; Department of Biomedical Sciences for Health, University of Milan, Milan, Italy; Department of Neurorehabilitation Sciences, Casa di Cura del Policlinico, Milan, Italy; Neuromuscular and Rare Diseases Unit, Department of Neuroscience, Fondazione IRCCS Ca’ Granda Ospedale Maggiore Policlinico, Milan, Italy; Department of Life Sciences, University of Trieste, Trieste, Italy; UCL Genetics Institute, Department of Genetics, Evolution and Environment, University College London, UK

**Keywords:** alternative splicing, ALS-FTLD, IBM, muscle, TDP-43

## Abstract

TDP-43 aggregation and redistribution have been recognised as a hallmark of amyotrophic lateral sclerosis, frontotemporal dementia and other neurological disorders. While TDP-43 has been studied extensively in neuronal tissues, TDP-43 inclusions have also been described in the muscle of inclusion body myositis patients, highlighting the need to understand the role of TDP-43 beyond the central nervous system. Using RNA-seq we performed the first direct comparison of TDP-43-mediated transcription and alternative splicing in muscle (C2C12) and neuronal (NSC34) mouse cells. Our results clearly show that TDP-43 displays a tissue-characteristic behaviour targeting unique transcripts in each cell type. This is not due to variable transcript abundance but rather due to cell-specific expression of RNA-binding proteins, which influences TDP-43 performance. Among splicing events commonly dysregulated in both cell lines, we identified some that are TDP-43-dependent also in human cells and show that inclusion levels of these alternative exons appear to be differentially altered in affected tissues of FTLD and IBM patients. We therefore propose that TDP-43 dysfunction, reflected in aberrant splicing, contributes to disease development but it does so in a tissue- and disease-specific manner.

## INTRODUCTION

TDP-43, a protein encoded by the *TARDBP* gene, is a ubiquitously expressed member of hnRNP family able to bind DNA and RNA that participates in various steps of mRNA metabolism including transcription, pre-mRNA splicing, miRNA generation, regulation of mRNA stability, nucleo-cytoplasmic transport and translation (Birsa *et al*, 2020; Budini & Buratti, 2011; Ederle & Dormann, 2017; Buratti & Baralle, 2012). TDP-43 was initially described as the major component of cytoplasmic inclusions formed in motor neurons of patients suffering from amyotrophic lateral sclerosis (ALS) and frontotemporal dementia (FTLD) despite the fact that mutations in *TARDBP* gene only account for a small subset of those cases (Arai *et al*, 2006; Buratti, 2015; Neumann *et al*, 2006). However, TDP-43 aggregates have as well been found in skeletal muscles of patients with inclusion body myositis (IBM) (Salajegheh *et al*, 2009; Weihl *et al*, 2008), oculopharyngeal muscular dystrophy (OPMD) (Yamashita *et al*, 2013) and limb girdle muscular dystrophy type 2a (LGMD2a) (Harms *et al*, 2012) suggesting that TDP-43 aggregation may play a prominent pathological role also in muscle tissue. Accordingly, TDP-43 myogranules have been shown to provide essential functions during skeletal muscle development and regeneration, both in mouse and human (Vogler *et al*, 2018). Despite ubiquitous expression of TDP-43, however, most studies investigating this protein have focused on its role in the central nervous system. Nonetheless, given its importance of TDP-43, both in muscle development and potentially in the pathogenesis of numerous myopathies, we systematically investigate functions elicited by TDP-43 in muscle (C2C12) and neuronal (NSC34) mouse cells in parallel.

Performing such a comparison is particularly interesting as these two cell environments display tissue-characteristic features, like for example: distinct post-translational modifications (PTMs) and cleavage products of TDP-43 described in muscles and neurons (Buratti, 2018), muscle-characteristic localization of TDP-43 in space and time (Vogler *et al*, 2018), cell-type-specific milieu of TDP-43 binding partners (Mele *et al*, 2015), and differential expression of RNA binding proteins (RBPs) controlling common mRNA targets (Appocher *et al*, 2017; Cappelli *et al*, 2018). It is important to note that all these differences occur in a context of highly variable transcriptome between tissues including non-coding transcripts (Cabili *et al*, 2011; Jiang *et al*, 2016; Ludwig *et al*, 2016). Therefore, TDP-43 might likely elicit tissue characteristic functions by targeting unique subsets of transcripts, which encode proteins participating in tissue-specific cellular pathways and provide crucial structural and functional features of a cell. The consequences of TDP-43 dysfunction in muscles could thus possibly differ from those that have so far been described in the central nervous tissue (Polymenidou *et al*, 2011; Tollervey *et al*, 2011).

In the last decade, high throughput methodologies have shifted the focus from characterization of individual events towards less biased global approaches, setting the ground for a systematic comparison of TDP-43 targeted RNAs across tissues and conditions. However, the overlap of TDP-43-controlled events identified by earlier studies is rather poor. It probably reflects the variation in technical approaches (microarrays, RNA-seq, CLIP-seq) and models employed in those studies: mouse brain (Polymenidou *et al*, 2011), human-post mortem brain samples (Tollervey *et al*, 2011; Prudencio *et al*, 2020), human neuroblastoma cell line SH-SY5Y (Fiesel *et al*, 2012; Tollervey *et al*, 2011), HEK-293 (De Conti *et al*, 2015; Prpar Mihevc *et al*, 2016), Hela (Prudencio *et al*, 2012). A clearer understanding of the extent to which TDP-43-mediated events are conserved between mouse and human is still lacking, yet it is a crucial point that should be addressed in future as it will allow better comparisons of human and mouse models of disease.

To finally address this issue in a systematic manner, we have identified subsets of unique cell-type-specific mRNA targets, as well as commonly regulated mRNAs, the tight regulation of which might underlie functions crucial for cell survival. More specifically, we have further explored splicing events that commonly occur in C2C12 and NSC34 cells and are additionally conserved in humans. We finally show that inclusion of common mouse-human TDP-43-regulated alternative exons is indeed altered in skeletal muscles of IBM patients and different brain regions of ALS and FTLD patients with reported TDP-43 pathology.

## RESULTS

### TDP-43 expression is similar in C2C12 and NSC34 cells

To start comparing the functions of TDP-43 in cells of muscular and neuronal origin, we used the most commonly employed mouse cell lines representing skeletal muscle (C2C12) and motor neurons (NSC34). They have been previously used to study TDP-43-associated neurodegeneration as well as the role of TDP-43 in muscle development (Budini *et al*, 2015; Colombrita *et al*, 2009; Militello *et al*, 2018; Vogler *et al*, 2018). We first assessed protein levels of endogenous TDP-43 in untreated cells (**Fig 1A**). Although in mature mouse tissues TDP-43 expression was reported to be higher in the brain compared to quadriceps muscle (Jeong *et al*, 2017), we noted no difference in the amount of total TDP-43 protein between undifferentiated C2C12 and NSC34 cells (**Fig 1A**), nor in the expression of TDP-43 at the RNA level of siLUC-transfected cells (**Fig 1B**).

**Figure 1.**
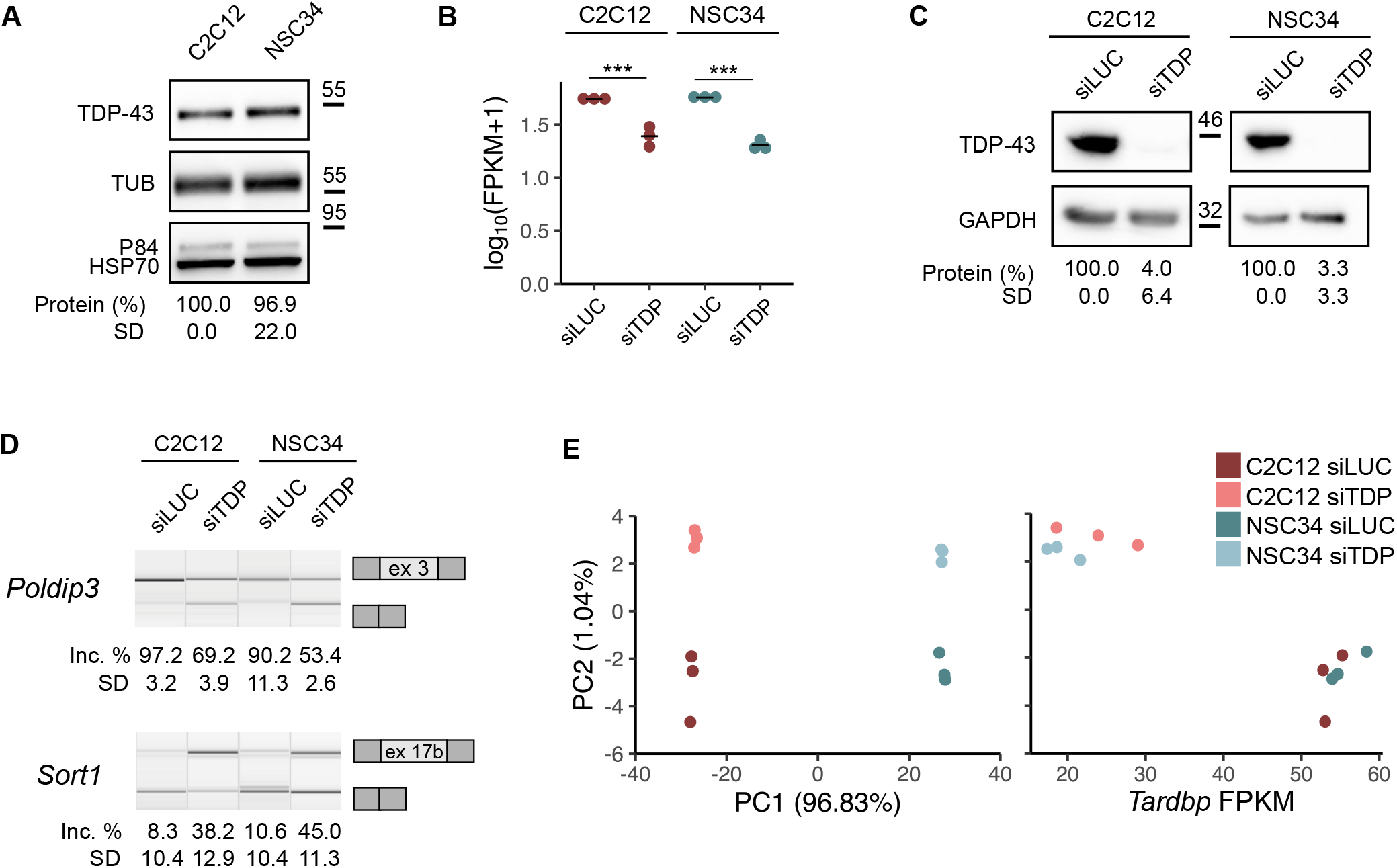
TDP-43 expression and functional consequences of TDP-43 silencing in C2C12 and NSC34 cells. **A** Western blot shows similar expression of endogenous TDP-43 in C2C12 and NSC34 cells. The amount of TDP-43 was normalized to the sum of peak intensities of three loading controls (tubulin, HSP70 and P84) (n = 3 replicates per group). **B** Expression levels of *Tardbp* in TDP-43-silenced C2C12 and NSC34 and corresponding controls assessed by RNA-sew plotted as log_10_-transformed FPKM values show TDP-43 was depleted (on the mRNA level) to the same extent in both cell lines (n = 3 replicates per group). p_adj_ = 1.6 · 10^-18^ for C2C12 and p_adj_ = 2.3·10^-72^ for NSC34. p-values were generated using Wald test and Bejamini-Hochberg multiple testing correction (Love *et al*, 2014). **C** Western blot shows the reduction of TDP-43 in C2C12 and NSC34 cells upon siTDP transfection. siLUC-transfected cells were used as a control. TDP-43 expression was normalized against GAPDH (n = 3 replicates per group). **D** TDP-43 depletion led to altered splicing of *Poldip3* and *Sort1*. Semi quantitative RT-PCRs conducted in TDP-43-silenced samples and corresponding controls are shown along with the quantification of splicing changes (% of alternative exon inclusion). The number of the alternative exon is given below (see the exact transcript numbers in **Appendix Table S1**, n = 3 replicates per group). **E** PCA plot visualizes distances between siLUC- and siTDP-transfected C2C12 and NSC34 cells based on FPKM of all genes obtained by RNA-seq (left). Variation in the PC2 is explained by the presence/absence of TDP-43 (right).

TDP-43 was silenced to a similar extent in both cell lines (**Fig 1B**) and reduction of the protein was confirmed by western blot (**Fig 1C**). TDP-43 loss functionally reflected in altered splicing of the two well characterized target transcripts *Poldip3* and *Sort1* (**Fig 1D**) (Fiesel *et al*, 2012; Mohagheghi *et al*, 2016; Prudencio *et al*, 2012; Shiga *et al*, 2012). To explore transcriptome-wide effects of TDP-43 downregulation, we then performed deep RNA-seq analysis on polyadenylated mRNA extracted from TDP-43 depleted cells. Both cell lines displayed a characteristic transcriptional signature as revealed by PCA (PC1), whereas the effect of TDP-43 knockdown explained a smaller portion of the variation between samples (PC2) (**Fig 1E**). This result suggests that TDP-43 silencing promotes transcriptional alterations in C2C12 and NSC34 based on the tissue-characteristic transcriptional profile.

### mRNAs dysregulation following TDP-43 reduction in C2C12 and NSC34 cells is cell-type specific

Tissues vary substantially in transcription levels of individual genes and splice isoforms they express, and these differences underlie specific biological characteristics and functions. To examine the effect of TDP-43 loss on expression levels (differential gene expression, DEG) in the two cell lines, we separately normalized reads of C2C12 and NSC34 datasets and obtained 4019 transcripts, expression levels of which were subject to TDP-43 regulation. At p_adj_ < 0.05, we detected a very similar number of DEG in C2C12 and NSC34 (2325 and 2324, respectively), with 630 (15.7%) transcripts being commonly dysregulated in both cell lines (**Fig 2A**). Surprisingly enough, the small overlap could not be explained by the fact that some genes are expressed in a tissue-specific manner (i.e., muscle characteristic genes are not transcribed in neuronal cells and *vice versa*), as the overlap between TDP-43 targets remained small (19.3%) even if we only considered genes expressed in both cell lines (FPKM in both cell lines > 0.5) (**Fig 2B**). However, our data indicated that TDP-43 targets regulated in a cell-type-specific fashion are highly expressed in one cell type but not in the other. On average, C2C12-specific TDP-43-regulated mRNAs show higher expression in C2C12 than in NSC34 cells, and *vice versa* (**Appendix Fig S1A**).

**Figure 2.**
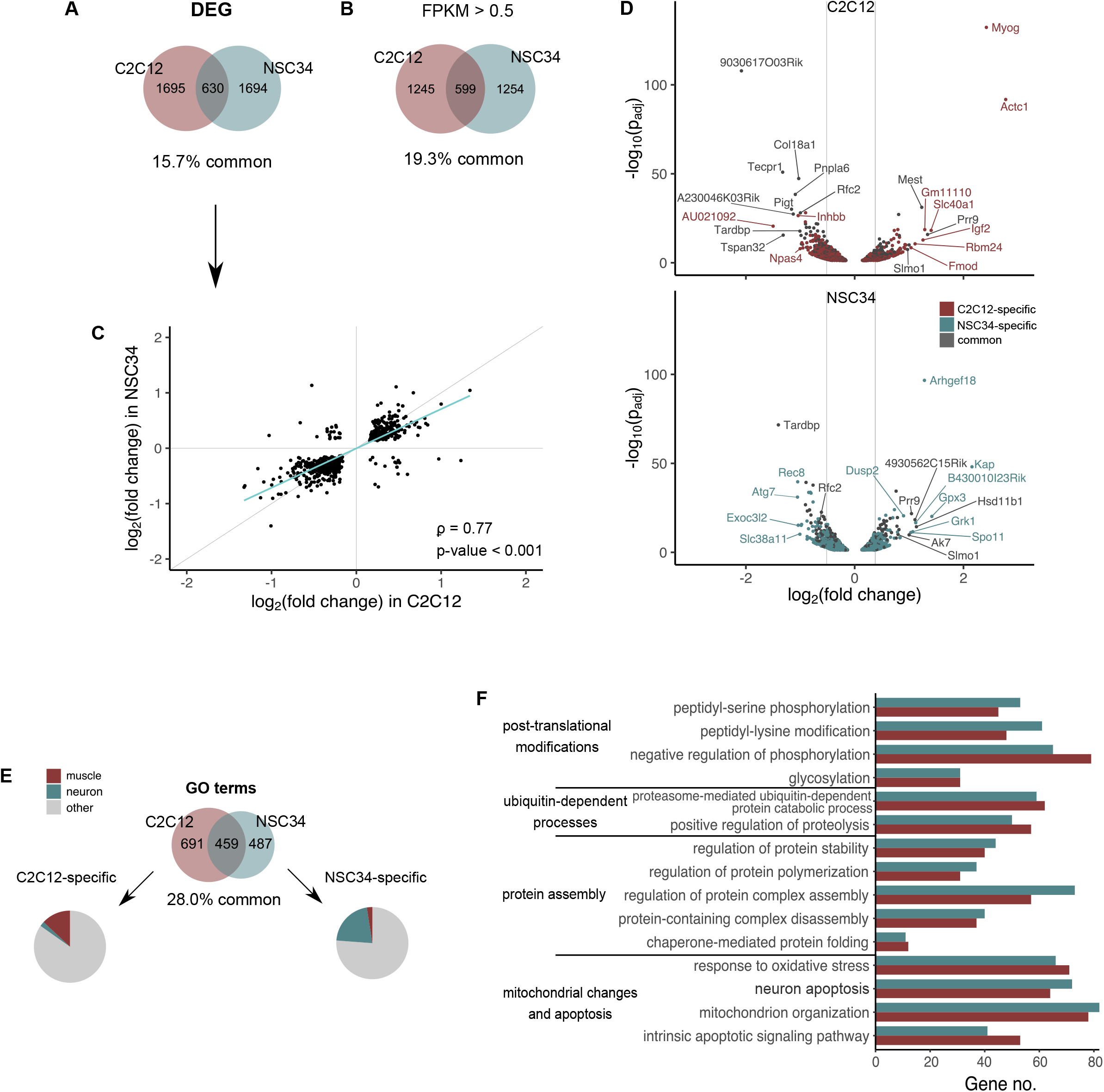
TDP-43 mediates transcription levels of different mRNAs in C2C12 and NSC34 cells. **A** Venn diagram shows the number of TDP-43-regulated transcripts identified in C2C12 and NSC34 cells exclusively (1695 and 1694, respectively), along with those that are commonly regulated by TDP-43 in both cell types (630). Transcripts with p_adj_ < 0.05 were considered as differentially expressed irrespective their log2 fold change. **B** The Venn diagram shows the overlap (599 transcripts, 19.3%) of TDP-43-regulated DEG identified in C2C12 and NSC34 cell line (as in (**A**)), considering only transcripts expressed in both cell lines (FPKM in both cell lines > 0.5). Transcripts with p_adj_ < 0.05 were considered as differentially expressed irrespective their log2 fold change. **C** Transcription changes of common targets ((**A**), 630) are plotted by their log2 fold change values in C2C12 and NSC34 (Spearman’s ρ = 0.77, p-value < 2.2 ·10^-16^). Grey line represents y = x and the blue line represents the fitted regression. **D** TDP-43-mediated transcription changes in C2C12 and NSC34 represented as volcano plots. C2C12- and NSC34-specific targets are shown in red and blue, respectively, while common targets are plotted as grey dots. Vertical lines indicate fold changes of 0.7 (30% increase) and 1.3 (30% decrease). Best hits are labelled with gene names. **E** The Venn diagram shows the number of cell-type-specific and overlapping GO terms enriched by DEGs identified in C2C12 or NSC34 cells. GO terms (category: biological process) were grouped based on their names as those implying muscle- (red) or neuron-related features (blue). **F** Representative GO terms (category: biological process) commonly enriched by DEGs in C2C12 and NSC34 cells suggesting pathological abnormalities described in neurodegenerative and myodegenerative disease (hand curated).

It has previously been proposed that TDP-43 binding is needed to sustain pre-mRNA levels and that mRNA downregulation would be a direct consequence of TDP-43 loss, while mRNA upregulation was explained by indirect effects (Polymenidou *et al*, 2011). In our datasets (**Fig 2A**), the number of downregulated genes slightly outnumbered genes that were upregulated following TDP-43 depletion (**Appendix Fig S1B**), however, the overlap was very similar, irrespective the direction of the change (14.0% and 15.0% for upregulated and downregulated transcripts, respectively). Comparing the extent of expression changes of commonly regulated transcripts (630) induced by TDP-43 reduction, we saw a positive correlation (ϕ = 0.77, p-value < 0.001) between the two cell lines, with a trend towards larger alternations in C2C12 (**Fig 2C**). Of note, there were few mRNAs whose expression was altered in the opposite direction in the two cell lines, indicating that TDP-43 loss can elicit contrary effects (*loss-of-function* vs. *gain-of-function*) depending on the cellular environment. Looking at individual target transcripts (**Fig 2D**, **Appendix Fig S1C and D**), we hypothesized that the biggest transcriptional changes induced by TDP-43 loss occurred in highly expressed genes. However, plotting the size of the change (log2 fold change) against background expression levels (FPKM in siLUC transfected cells) of all DEGs revealed that there is in fact no correlation between the two (**Appendix Fig S1E**).

Taken together, these results support the idea that unique sets of transcripts controlled by TDP-43 in each cell type can only partially be explained by variable expression levels of cell-type-characteristic mRNAs across tissues. Factors other than expression levels as such thus influence TDP-43 function that seems to be tissue-specific. At the sequence level, in fact, TDP-43-regulated mRNAs detected in C2C12 or NSC34 appear to be equally well conserved across species (**Appendix Fig S1F**).

### Commonly enriched processes implicated in neurodegenerative and myodegenerative disease

In the mouse brain, TDP-43 has been shown crucial for maintenance of mRNAs that encode proteins involved in synaptic activity (Polymenidou *et al*, 2011). To elucidate which cellular processes might be controlled by TDP-43 in cells of muscle and neuronal origin, we conducted enrichment analysis of genes differentially expressed in C2C12 (2325) and NSC34 (2324) (**Fig 2E**). Among C2C12 enriched GO terms, we found those directly associated with muscle characteristic features like *striated muscle development* or *muscle cell migration*, in line with results highlighting the importance of TDP-43 in skeletal muscle formation and regeneration (Militello *et al*, 2018; Vogler *et al*, 2018). On the other hand, a great portion of neuronal processes like *vesicle-mediated transport in synapse* or *regulation of postsynaptic membrane neurotransmitters* appeared to be affected by TDP-43 loss in NSC34 cells.

While the percentage of overlapping DEG was only 15.7%, by GO categories, almost a third of all biological processes (28%) enriched in C2C12 or NSC34 DEGs (**Fig 2E**) was commonly dysregulated upon TDP-43 depletion in both cell lines. Given that currently proposed picture of pathological processes implicated in myopathies bears several similarities with neurodegenerative disease (Askanas *et al*, 2015, 2012; Weihl *et al*, 2008), we investigated commonly enriched GO terms to see, if any of them could detect abnormalities previously described in the above-mentioned diseases. Significant GO terms enriched by DEG in both C2C12 and NSC34 (**Fig 2F**) suggest that some common TDP-43-mediated mechanisms might contribute to development of TDP-43-proteinopathies in both muscle and neuronal tissues. Pathomechanisms include aberrant protein accumulation (i.e., ubiquitin, amyloid β, α-synuclein, phosphorylated τ and TDP-43), post-translational modifications of deposited proteins (phosphorylation, ubiquitination, acetylation, sumoylation), defects in protein disposal (26S proteasome and autophagy) and mitochondrial abnormalities. However, while there was a greater overlap between biological response to TDP-43 depletion (GO: biological process), the specific differentially expressed transcripts in common terms were remarkably different between C2C12 and NSC34 (**Appendix Fig S1G**). This implies that TDP-43 can influence similar biological processes in both muscles and neurons, but it does so by mediating expression levels of genes encoding for distinct proteins that participate in those pathways.

### TDP-43-mediated splicing is more pronounced in NSC34 cells

Along with mRNA depletion, aberrant pre-mRNA splicing has been described to contribute to neuronal vulnerability as a consequence of pathologic TDP-43 behaviour (Arnold *et al*, 2013; Polymenidou *et al*, 2011; Tollervey *et al*, 2011). Yet, little is understood about how TDP-43 dysfunction affects pre-mRNA splicing in tissues beyond the central nervous system. In this work, we systematically compared alternative splicing (AS) alterations following TDP-43 reduction in C2C12 and NSC34 cells. As expected, a considerably lower number of splicing events was detected in C2C12 than in NSC34 cells (730 and 1270, respectively) at FDR of 0.01 (**Fig 3A**), which held true for events of any classical AS category (i.e., SE, MXE, RI, A3’SS, A5’SS) (**Fig 3B**). Neuronal and muscular targets did not vary with regard to event type proportion (**Appendix Fig S2A**); length of cassette exons (**Appendix Fig S2B**); the ratio between inclusion/exclusion events (**Appendix Fig S2C**); or percentage of frame-conserving events (**Appendix Fig S2D**). Interestingly enough, alternative sequences regulated by TDP-43 in the neuronal cell line seem to be more conserved across species than TDP-43-regulated sequences in muscle cell line (**Appendix Fig S2E**). This holds true particularly for cassette exons (**Appendix Fig S2F**), which represent the most frequent event type detected by our pipeline (**Appendix Fig S2A**).

**Figure 3.**
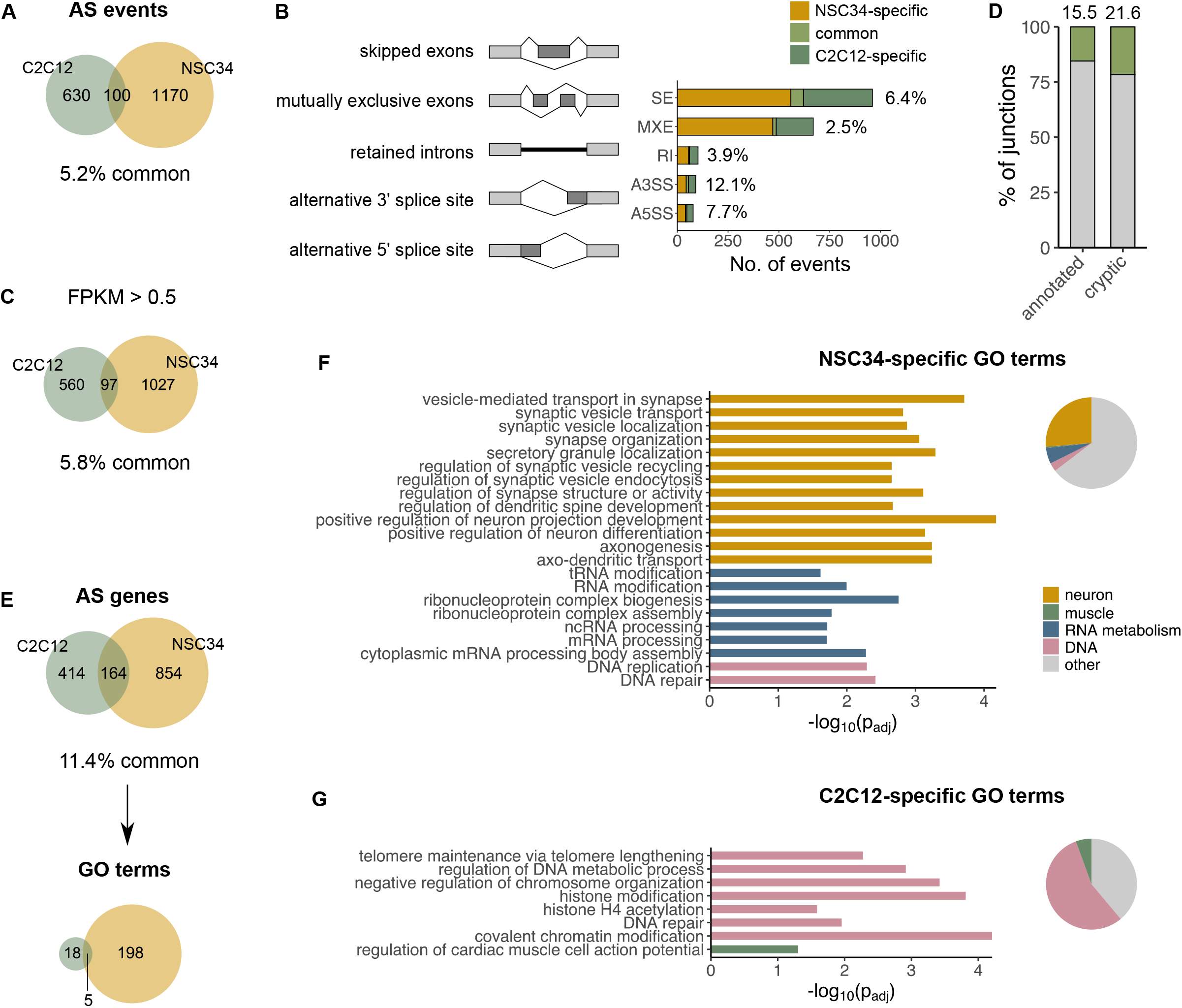
TDP-43-regulated splicing changes show cell-type specificity. **A** Venn diagram shows the total number of AS events (detected by rMATS at FDR < 0.01) induced by TDP-43 depletion in C2C12 and NSC34 specifically (630 and 1170, respectively), together with those commonly detected in both cell lines (100). **B** The number of annotated AS events (**A**) visualized by event type. SE - exon skipping, MXE - mutually exclusive exons, RI - intron retention, A3’SS and A5’SS - alternative 3’ or 5’ splice site. The percentage of overlapping AS events is reported on the plot. **C** Venn diagram shows the total number of AS events (detected by rMATS as in (**A**)) occurring in transcripts, which are expressed in both cell lines (FPKM in both cell lines > 0.5). **D** The percentage of common (green) and cell-type-specific (grey) TDP-43-dependent splicing events detected in C2C12 and NSC34 as assessed by MAJIQ (in contrast to (**A)-(C)** and (**E)- (G)**, where rMATS was used). **E** Venn diagrams show the number of alternatively spliced transcripts (as detected by rMATS, FDR < 0.01) in C2C12 and NSC34 cells together with GO terms (category: biological process, p_adj_ < 0.05) enriched in AS genes detected in each cell line. **F** GO terms uniquely enriched in NSC34 (198) imply on deregulation of neuronal processes, mRNA metabolism and DNA biology in NSC34 cells (representative GO terms are shown on the plot). **G** GO terms uniquely enriched in C2C12 (18) suggest involvement of TDP-43-regulated AS genes in DNA-modifying processes (representative GO terms are shown on the plot).

This observation that TDP-43 regulates more events in NSC34 cells might reflect the importance of alternative splicing as a regulatory mechanism in neurons and support the existence of a distinct splicing program in neuronal tissues, as already suggested by others (Irimia *et al*, 2014; Mele *et al*, 2015; Yeo *et al*, 2004). Moreover, very few AS events (on average 5.2%) appear to be commonly regulated by TDP-43 in both cell types, with the percentage of overlapping AS events being small (5.8%) even when we only considered AS in transcripts commonly expressed in both cell lines (FPKM > 0.5) (**Fig 3C**) or when we used a less stringent overlap threshold (**Appendix Fig S2G**).

Jeong et al. (Jeong *et al*, 2017) have previously reported that TDP-43’s repression of cryptic exons is tissue-specific. This posed a question whether annotated TDP-43-controlled events (**Fig 3A** and **3B**) display more or less tissue-variation compared to TDP-43-repressed cryptic exons. As rMATS, the splicing tool used to identify annotated AS events, is not capable of identifying non-canonical splicing we used a separate analysis tool, MAJIQ (Green *et al*, 2018), that allows quantification of both, novel (cryptic) and regular (annotated) AS events. MAJIQ and rMATS quantify in separate ways (Mehmood *et al*, 2020), thus comparable results are only produced (junctions or AS events, respectively) when the same pipeline is applied. Using MAJIQ, we show that the percentage of commonly detected cryptic splicing is in fact bigger than that of commonly detected classical AS events (**Fig 3D**) (21.6% and 15.5%, respectively), implying that TDP-43 displays tissue-specific behaviour in cryptic repression but even more so in control of classical alternative exons.

### Alternatively spliced TDP-43 targets are implicated in neuronal functions and DNA-related processes

We further employed GO analysis to see whether genes with TDP-43-regulated splicing identified in C2C12 and NSC34 form interconnected networks and if TDP-43 can, by mediating AS, influence particular biological processes in each cell type. Since the number of C2C12 AS genes entering GO analysis (578) was considerably lower than that of NSC34 genes (1018), the analysis resulted in fewer GO terms found to be enriched in C2C12 compared to many in NSC34 (23 and 203, respectively) (**Fig 3E**). As expected, GO terms enriched in NSC34 cells exclusively suggest that in these cells, alternatively spliced mRNA predominantly encode for proteins implicated in processes taking place in the nervous system (e.g., *axonogenesis, regulation of neuron differentiation*) (**Fig 3F**). This is in line with earlier studies, which demonstrated that in human neuroblastoma cells SH-SY5Y TDP-43-dependent splice isoforms encode for proteins regulating neuronal development and those involved in neurodegenerative disease (Tollervey *et al*, 2011).

On the other hand, GO terms (56%) enriched in C2C12 cells exclusively (**Fig 3G**) suggested involvement of AS genes in DNA-related processes (e.g., *covalent chromatin modification* or *regulation of chromosome organization*), while only one implied a muscle characteristic feature (i.e., *regulation of cardiac muscle cell action potential*). As we thought this observation might be biased due to the low number of GO terms detected in C2C12 (18), we repeated enrichment analysis, this time using a more relaxed threshold (non-corrected p-value < 0.01 instead of FDR < 0.01) on AS genes that would enter GO analysis. However, even among 45 enriched GO terms obtained using less stringent threshold, DNA-related processes comprised more than a third of all GO terms (36%, **Appendix Fig S2H**), which was not the case for NSC34 cells.

### Different RBPs are expressed in NSC34 and C2C12 cells

The observation that TDP-43 loss elicits a tissue-characteristic response did not come as a surprise, as RNA binding proteins (RBPs) other than TDP-43 might be differentially expressed in these cells. Inspecting expression levels of some RNA-binding proteins (Mele *et al*, 2015), which either directly interact with TDP-43 (Freibaum *et al*, 2010) or influence processing of its target transcripts (Cappelli *et al*, 2018; Lagier-Tourenne *et al*, 2012; Mohagheghi *et al*, 2016), we saw a higher average expression of RBPs in neuronal NSC34 cells (**Fig 4A**) in line with previous observations (Mele *et al*, 2015). Their joint functions in coordinating mRNA processing might underlie a more complex splicing regulation that is unique for neuronal tissues and explain why TDP-43-regulated splicing is more frequent in NSC34 than in C2C12 cells (**Fig 3A**). The two cell types clearly express a distinct array of RBPs (**Fig 4B**), while transcription levels of some are additionally affected by TDP-43 depletion (**Fig 4C**).

**Figure 4.**
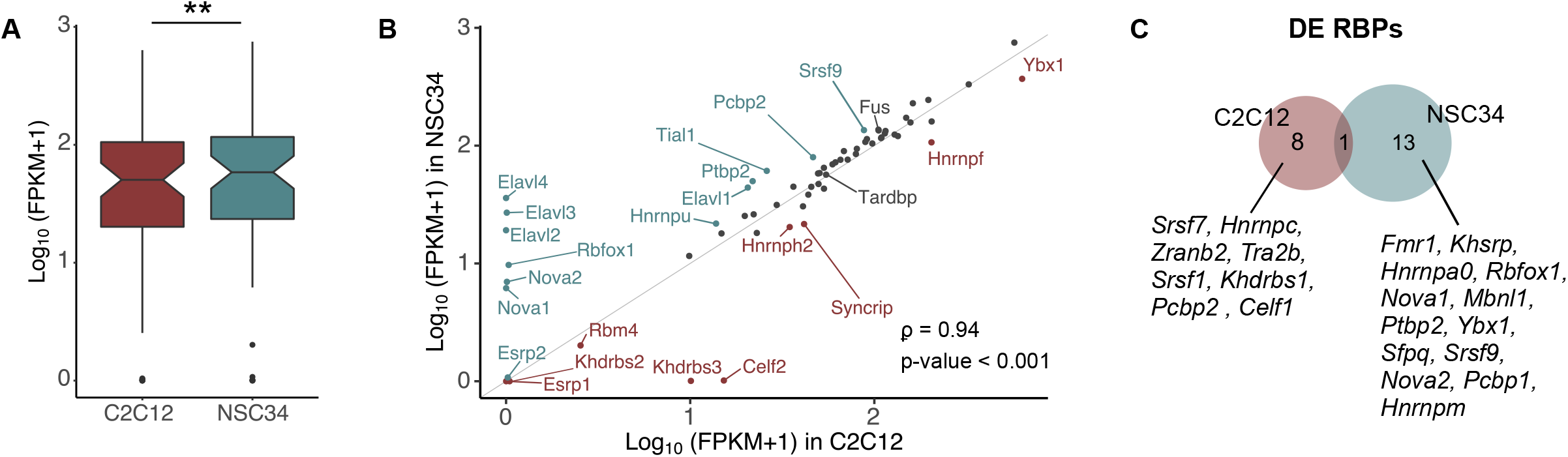
Expression of RNA-binding proteins in C2C12 and NSC34 cells. **A** Boxplot shows that NSC34 cells on average display higher expression of 63 RNA-binding proteins compared (Mele *et al*, 2015) compared to C2C12 cells (p-value = 0.0028). Average expression levels are plotted as log_10_-transformed FPKM values of all 63 transcripts and p-value was generated using Wilcoxon signed-rank test. **B** Expression of 63 RBPs (plotted as log_10_-transformed FPKM values) in C2C12 and NSC34 cells (Spearman’s ρ = 0.94; p-value < 2.2 · 10^-16^). Those with higher expression in one cell line than another (> 150%) are shown in red (C2C12) or blue (NSC34). Grey line represents y = x. **C** Venn diagram shows RBPs the expression of which changes following TDP-43 reduction. The overlapping event is downregulation of *Tardbp*.

### Common TDP-43 splicing targets detected in C2C12 and NSC34

Previous studies have already disclosed lists of transcripts, whose splicing is affected by TDP-43 removal or dysfunction (Colombrita *et al*, 2009; De Conti *et al*, 2015; Lagier-Tourenne *et al*, 2012; Tollervey *et al*, 2011). Yet, the reproducibility of target identification is rather poor, possibly due to differences in methodological approaches, low conservation of TDP-43 targets across species (Colombrita *et al*, 2009), and, as we show, the unique function TDP-43 elicits in each tissue or cell type. The most consistently reported TDP-43-regulated splicing event across studies and conditions is skipping exon 3 within *Poldip3/POLDIP3* mRNA (both mouse and human) (Fiesel *et al*, 2012). This being so, inclusion level (percent spliced in,ΔPSI) of *Poldip3* exon 3 often serves as a readout of TDP-43 functionality (Cortese *et al*, 2018; Klim *et al*, 2019; Roczniak-Ferguson & Ferguson, 2020). In search of new splicing events that would, similarly to *Poldip3/POLDIP3*, show high reproducibility across experimental settings, we chose mRNAs that underwent the biggest shift in TDP-43-dependent exon inclusion and whose isoform proportion was altered in both C2C12 and NSC34 cells. The isoform switch of these targets was validated using isoform-sensitive semi-quantitative RT-PCR (**Fig 5A**).

**Figure 5.**
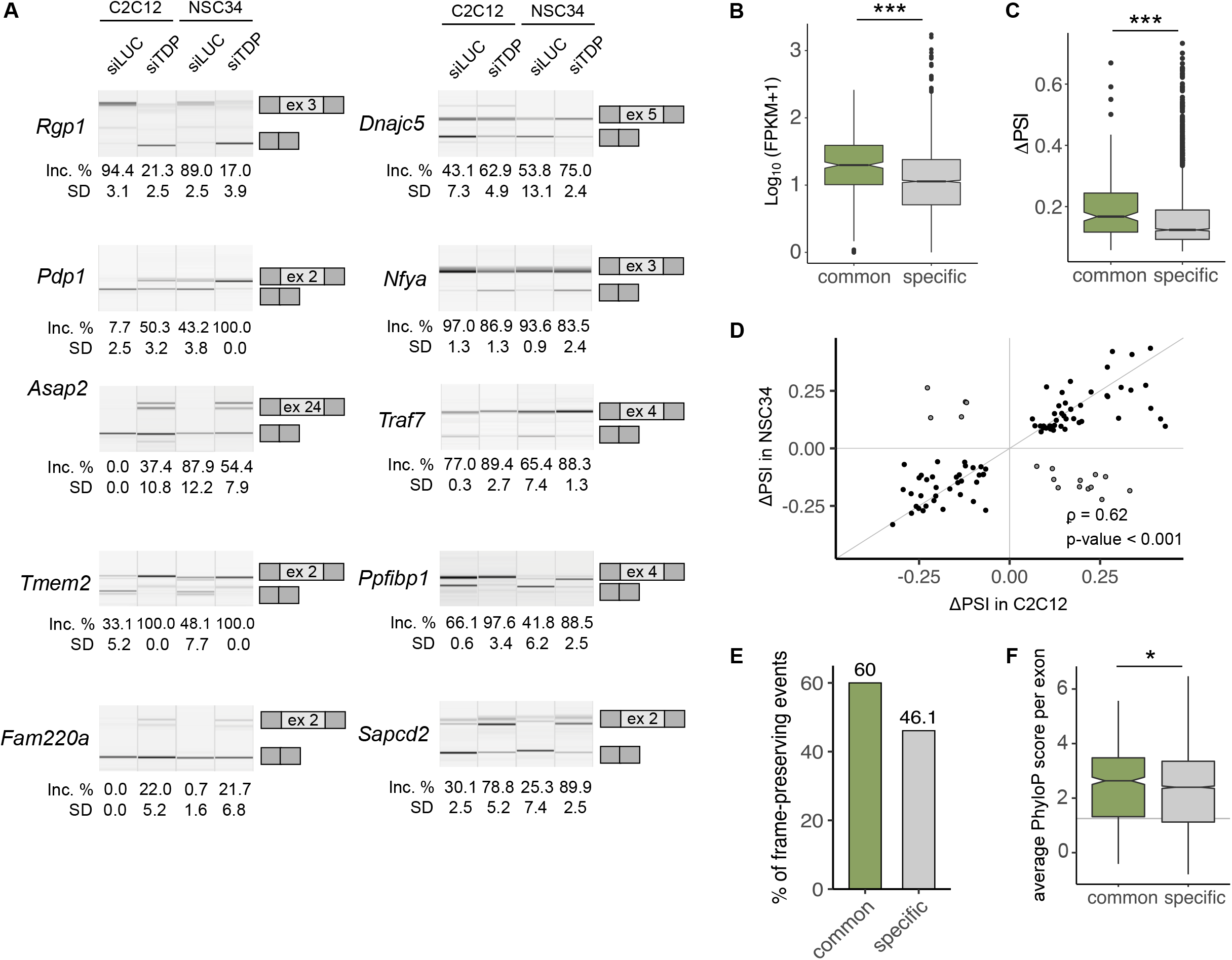
Commonly regulated TDP-43 splicing targets are more often frame-conserving, display higher expression levels and undergo bigger changes in isoform proportion. **A** Validation of TDP-43 dependent splicing of 10 representative mRNA targets. Semi quantitative RT-PCR conducted in TDP-43-silenced samples and corresponding controls is shown along with the quantification of splicing changes (% of alternative exon inclusion). The number of the alternative exon is shown in the scheme (see the exact transcript numbers in **Appendix Table S1**, n = 3 replicates per group). **B** Average expression expression levels of transcripts that are commonly spliced in both cell lines (164) or in one cell line exclusively (1268) is plotted as log_10_-transformed FPKM values (p-value < 2.2·10^-16^). **C** Absolute changes (ΔPSI) of overlapping splicing events (100) compared to those uniquely occurring in C2C12 or NSC34 (1800) (p-value = 1.0·10^-7^). p-values for **(B)** and **(C)** were generated by unpaired Wilcoxon rank sum test. **D** The correlation of splicing changes for commonly detected splicing events (100) plotted as ΔPSI in C2C12 and NSC34 (Spearman’s correlation coefficient ρ = 0.62, p-value = 4.1·10^-12^). **E** The percentage of frame-preserving AS events among those that commonly occur in both cell lines (100) and those regulated by TDP-43 in a cell-type specific manner (1800). **F** Average *per exon* PhyloP conservation scores plotted as box plots show TDP-43-regulated alternative sequences detected in both cell lines (634) are better conserved across species than those detected in one cell line exclusively (6019) (p-value = 0.02). p-value was generated using Wilcoxon rank sum test, the grey line represents the median of average PhyloP scores of all exons in the mouse genome.

Compared to cell-type-specific TDP-43 targets, commonly spliced transcripts on average show higher expression in C2C12 and NSC34 cells than transcripts alternatively spliced in a cell-type-specific manner (**Fig 5B**). Furthermore, commonly detected events display bigger splicing transitions (bigger ΔPSI) (**Fig 4C**). Most of the splicing changes detected in C2C12 and NSC34 occurred in the same direction (83%, ρ = 0.62, p-value < 0.001) (**Fig 5D**), meaning that for that subset of transcripts, TDP-43 exerts a similar function in cells of neuronal and muscular background. We observed a higher frequency of frame-preservation among splicing events found to be controlled by TDP-43 in both cell lines (**Fig 5E**) along with better conservation of common TDP-43-regulated sequences across species (**Fig 5F**).

Alternative splicing occurs co-transcriptionally and the two mechanisms have been known to influence one another in a coordinated manner (Kornblihtt et al., 2013). In our case, however, only a small portion of transcripts undergoing TDP-43-dependent splicing additionally showed altered overall transcript abundance (21.9% and 21.2% in C2C12 and NSC34, respectively) (**Appendix Fig S3A**). At least in C2C12 cells, transcripts whose splicing was affected by loss of TDP-43 more often decreased in abundance, which might be indicative of nonsense mediated decay (**Appendix Fig S3B**). Finally, KEGG analysis performed on sets of differentially expressed or alternatively spliced genes suggest that TDP-43 knockdown could influence a particular molecular pathway such as axon guidance through change in transcript levels (DEG) or by the means of alternative splicing (**Appendix Fig S3C**).

### Novel TDP-43-regulated splicing events conserved between mouse and human

While incorporation of exon 3 into mature *Poldip3* mRNA is regulated by TDP-43 in both mouse and human cells (Fiesel *et al*, 2012), most of TDP-43’s regulated splicing has shown to be highly species and even tissue-specific. We therefore investigated if any of commonly detected TDP-43 targets (**Fig 5A**) are (according to VastDB (Tapial *et al*, 2017)) predicted to have an orthologous event in humans. Some TDP-43-mediated events found in mouse (*Rgp1* exon 3, *Sapcd2* exon 2, *Fam220a* exon 2) do not even have a corresponding orthologous exon in humans. For those with putative AS orthology (i.e., the presence of orthologous alternative exon in both species), we tested whether alternative exons were subject to TDP-43 control also in human cells. We silenced TDP-43 in two human cell lines representing neuronal and muscular cells – human neuroblastoma SH-SY5Y and rhabdomyosarcoma RH-30 (Reber *et al*,2016) (**Fig 6A**), which resulted in exon skipping within *POLDIP3* (**Fig 6B**). Likewise, TDP-43 depletion led to enhanced inclusion of exon 19 in *PPFIBP1* and exon 23 of *ASAP2* but not exon 5 of *TRAF7* or exon 3 of *NFYA* (**Fig 6C**). Although one study reported a great portion of TDP-43-controlled exons in mouse to have prior evidence of alternative splicing in humans (Polymenidou *et al*, 2011) we still lack understanding to what extent TDP-43 regulation of mRNA processing is conserved between species. Exon orthology (as assessed by sequence similarity) could not be predictive of AS conservation since exon incorporation into mature mRNA depends on the exonic sequence but also on *cis*-regulatory motives and *trans*-acting factors (Barbosa-Morais *et al*, 2012; Gueroussov *et al*, 2015; Raj & Blencowe, 2015). In fact, iCLIP performed in SH-SY5Y cells (Tollervey *et al*, 2011) identified direct TDP-43-binding sites in a close proximity of alternatively spliced exons within *PPFIBP1* and *ASAP2*, while that was not the case for *TRAF7* and *NFYA* (**Fig 6D**). This finding suggests that alternative exons of *PPFIBP1* and *ASAP2* found to be regulated by TDP-43 in mouse and human cells are most likely controlled by TDP-43 in a direct fashion by its binding to regulatory sequences neighbouring splice sites.

**Figure 6.**
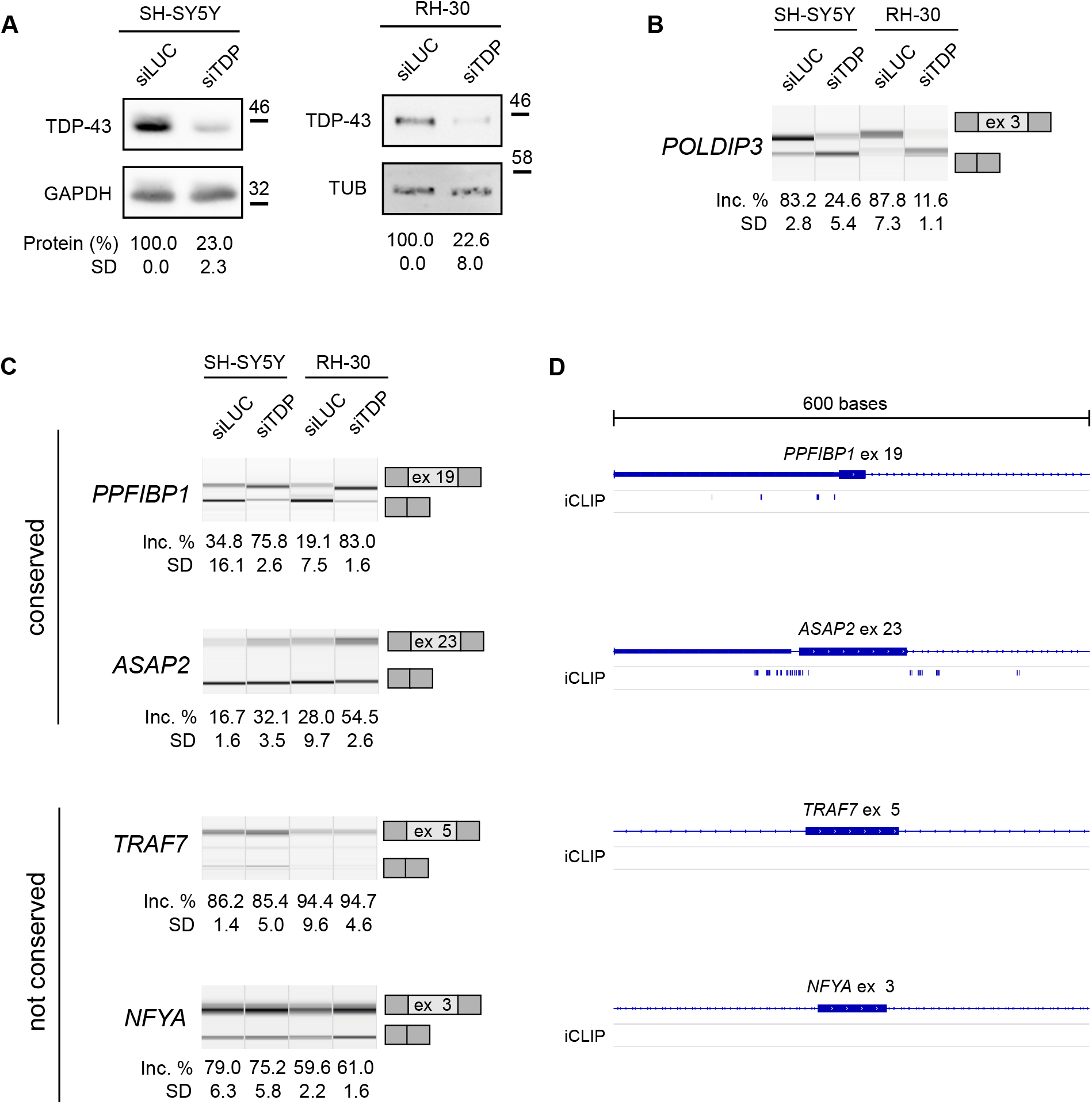
Alternative exons regulated by TDP-43 in mouse are subject to TDP-43 regulation in human cell lines or not. **A** Western blot shows efficient reduction of TDP-43 in SH-SY5Y and RH-30 cells upon siTDP transfection. The amount of TDP-43 was normalized against GAPDH or tubulin (n = 3 replicates per group). **B** TDP-43 depletion led to altered splicing of *POLDIP3*. Semi quantitative RT-PCR conducted in TDP-43-silenced samples and corresponding controls is shown along with the quantification of splicing changes (% of alternative exon inclusion). The number of the alternative exon is given below (n = 3 replicates per group). **C** Alternatively spliced exons regulated by TDP-43 in mouse cells are either subject to TDP-43 regulation in human cells (*PPFIBP1* exon 19 and *ASAP2* exon 23) or not (*TRAF7* exon 5 and *NFYA* exon 3). Semi quantitative RT-PCRs conducted in TDP-43-silenced samples and corresponding controls are shown along with the quantification of splicing changes (% of alternative exon inclusion). The number of the alternative exon is given in the scheme (see the exact transcript numbers in **Appendix Table S2**, n = 3 replicates per group). **D** Schematic representation of TDP-43 binding sites identified by iCLIP analysis in SH-SY5Y cells (Tollervey *et al*, 2011) in the vicinity of exons represented on panel **(C)**.

### Altered splicing patterns imply on TDP-43 dysfunction in FTLD and IBM patients

To explore if dysregulated alternative splicing could play a role in pathophysiology of TDP-43 proteinopathies, we measured inclusion levels of TDP-43-mediated alternative exons in IBM muscles (**Fig 7A**) as well as in pathological brain regions of ALS and FTLD cases with reported TDP-43 pathology (ALS-TDP and FTLD-TDP) (**Fig 7B** and **7C**). Since neuroanatomical regions markedly vary with regards to splice isoform expression (**Appendix Fig S4A**), we considered each brain region independently rather than analysing them together. Tissue-specific accumulation of truncated *STMN2*, which has recently been described as a very good clinical marker of TDP-43 impairment (Prudencio *et al*, 2020; Melamed *et al*, 2019; Klim *et al*, 2019), in fact occurs in brain areas previously known to be affected by TDP-43 pathology. We thus investigated TDP-43-controlled splicing in the spinal cord (lumbar and cervical, respectively) and the motor cortex of ALS cases (**Fig 7B**), whereas frontal and temporal cortices were the site of interest for FTLD patients (**Fig 7C**).

**Figure 7.**
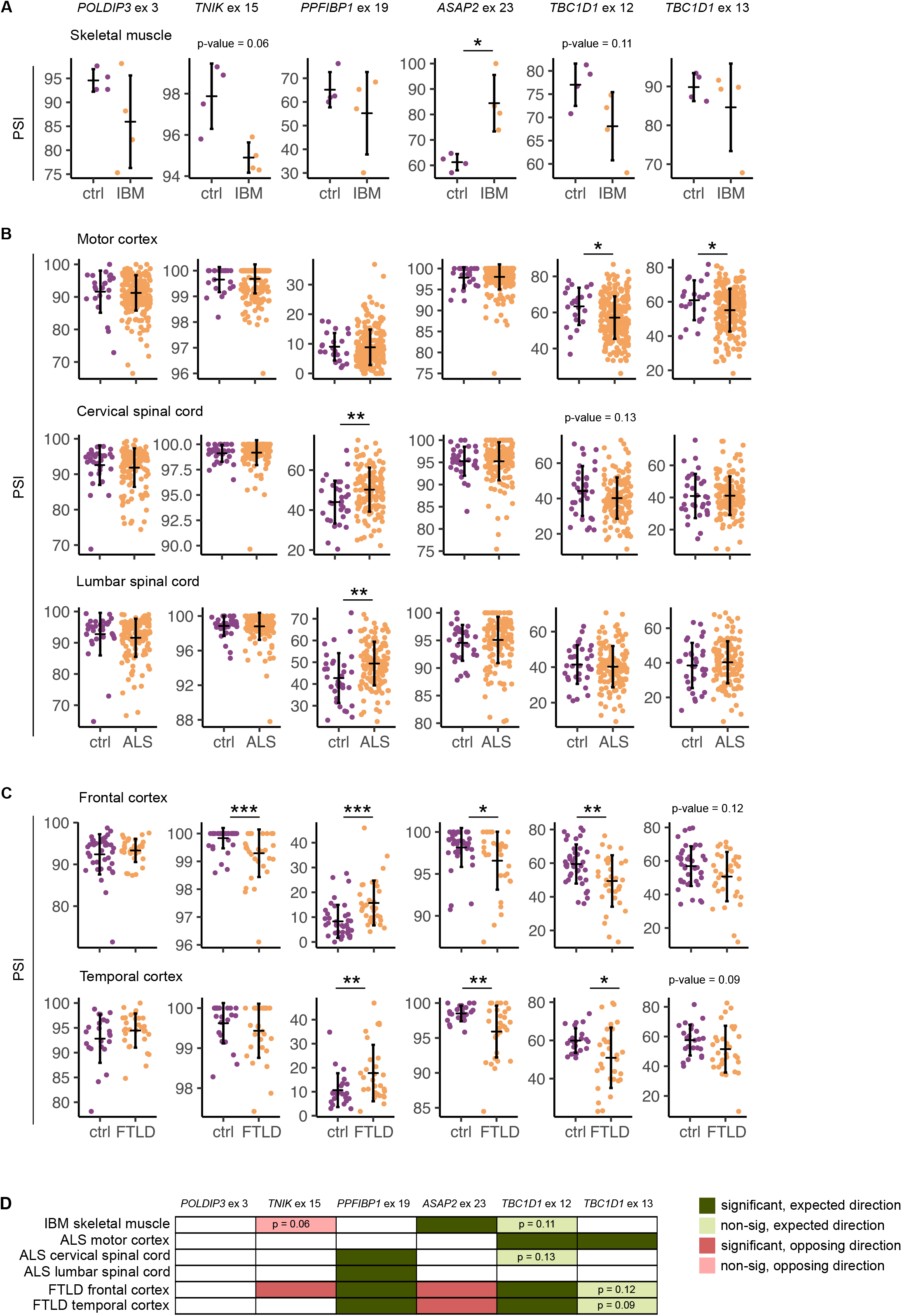
Inclusion of TDP-43-controlled exons is altered in TDP-43-proteinopathies. **A** Inclusion levels (PSI) of six alternative exons in skeletal muscle biopsies in IBM patients vs. healthy controls (n = 4 per group). **B** PSI of six alternative exons in different brain regions (motor cortex, lumbar spinal cord, cervical spinal cord) of ALS patients and healthy controls (n motor cortex: 223 ALS and 23 ctrl, n cervical spinal cord: 134 ALS and 32 ctrl, n lumbar spinal cord 136 ALS and 33 ctrl). **C** PSI of six alternative exons in frontal and temporal cortices of FTLD patients with reported TDP-43 pathology and healthy controls (n frontal cortex: 33 FTLD and 40 ctrl, n temporal cortex: 30 FTLD and 23 ctrl). **(A)**-**(C)** p-values were generated using Wilcoxon rank sum test. * p-value < 0.05, ** p-value < 0.01, *** p-value < 0.001. **D** Schematic summary of all splicing alterations **(A)**-**(C)** detected in skeletal muscles of IBM patients and across neuroanatomical regions of ALS and FTLD patients compared to healthy controls. Dark green marks significant changes, which occur in the same direction as in TDP-43-depleted SH-SY5Y and RH-30 cells (refers to **Fig 6C**); light green marks non-significant changes that occur in the expected direction (p-value is reported in the scheme); red marks significant changes occurring in the opposite direction relative to TDP-43-depleted SH-SY5Y and RH-30 cells; light red marks non-significant changes that occur in the opposite direction (p-value is reported in the scheme).

The splicing signature examined herein consisted of six TDP-43-regulated alternative exons: *POLDIP3* exon 3 is consistently detected as a TDP-43-regulated splicing event; exon 15 of *TNIK* has been previously described as TDP-43 target and was also detected in C2C12, SH-SY5Y and RH-30 cells (**Appendix Fig S4B** and **S4C**); exon 19 of *PPFIBP1* and exon 23 of *ASAP2* are newly identified TDP-43 targets conserved across species; exons 12 and 13 of *TBC1D1* are detected to be controlled by TDP-43 in C2C12 cells and are associated with muscle differentiation (Bland et al., 2010). The long *TBC1D1* isoform, which is dependent on TDP-43, appears to be crucial in mature tissues (Bland et al., 2010) but not in undifferentiated cells (as thus it was not detected in undifferentiated NSC34, SH-SY5Y and RH-30 cells (**Appendix Fig S4B** and **S4C**)). As TDP-43 binding sites were indeed found in the proximity of these alterative exons (**Appendix Fig S4D**), inclusion levels of exons 12 and 13 of *TBC1D1* gene were investigated in mature tissues coming from patients.

Interestingly, when we assessed inclusion levels of six TDP-43-controlled AS events in patient tissues, we got distinct patterns. For example, out of the six AS events, we only observed increased *ASAP2* exon 23 inclusion in IBM muscle relative to healthy controls (**Fig 7A**). In ALS cases, we detected significantly different inclusion of one exon (exon 19of *PPFIBP1*) in the lumbar and cervical spinal cord but not in the motor cortex, while in motor cortex, we saw enhanced skipping of both alternative exons within *TBC1D1* (**Fig 7B**). Surprisingly enough, FTLD appears to be the disease, in which splicing of six alternative exons is most heavily perturbed. Multiple TDP-43-targeted exons show significantly altered inclusion in patients, both in frontal and temporal cortices (**Fig 7C**). Apart from that, some non-significant changes clearly show a trend towards altered exon inclusion in patients.

At this point it is important to consider cell-type specific splicing activity of TDP-43 (**Fig 3A**), which makes it unlikely that upon TDP-43 malfunction splicing of the same transcripts would be altered across all cell types (**Fig 7D**). This being said, the scheme in **Fig 7D** summarizes splicing changes of six TDP-43-controlled exons detected in different tissues affected with TDP-43 pathology. The fact that splicing changes do not necessarily occur in the same direction as upon TDP-43 depletion in cell lines (as in the case of *ASAP2* and *TNIK*) again highlights the complexity of splicing control provided by TDP-43 that is generally acting within an interwoven network of splicing regulators. The same phenomenon (i.e., different directionality) was in fact observed when comparing the consequences TDP-43 depletion has on gene expression and alternative splicing *in vitro* using cell lines (**Fig 2C** and **5D**).

## DISCUSSION

TDP-43 inclusions represent the hallmark of ALS/FTLD (Arai *et al*, 2006; Neumann *et al*, 2006) and are frequently recognized as a secondary pathology in other neurodegenerative disease (Hasegawa *et al*, 2007; Higashi *et al*, 2007). In recent years, great progress has been made in explaining how potential *loss*- and *gain-of-function* mechanisms contribute to the pathogenesis observed in the brain and spinal cord (Budini *et al*, 2015; Cascella *et al*, 2016; Fratta *et al*, 2018). Nonetheless, a growing evidence of TDP-43 mis-localization and aggregation in tissues beyond the CNS has raised the possibility that TDP-43 dysfunction and consequently, impairment of RNA processing, might be deleterious for other tissues (Cortese *et al*, 2018, 2014).

To this date, cell- and tissue-characteristic molecular features of TDP-43 have seldom been investigated in parallel. Considering recent attention that TDP-43 has received in IBM and related pathologies (Harms *et al*, 2012; Salajegheh *et al*, 2009; Weihl *et al*, 2008; Yamashita *et al*, 2019), we therefore sought to fill this gap. The purpose of our study has been to further explicate the role of TDP-43 in different tissues to better understand its involvement in pathogenesis in cell types other than neurons, and to set the ground for development of potential therapeutic or biomarker strategies that focus on shared or specific disease mechanisms. We thus aimed to model *loss-of-function* effect in skeletal muscles vs. neurons and to focus on TDP-43-controlled alternative splicing (AS) events, as this is one of the best characterized features of this protein to date.

Although protein levels of TDP-43 itself are not different between C2C12 and NSC34, there is a tissue-characteristic expression of other RNA-binding proteins (e.g., those from *Elavl, Nova* and *Celf* families) that, like TDP-43, mediate RNA-related processes in a coordinated fashion. This result, together with differences in tissue-specific gene transcription levels, can presumably explain why there is little consistency across studies in identifying TDP-43-targeted transcripts (Buratti *et al*, 2013) and it clearly outlines the importance of cellular context in shaping the functional role of TDP-43. With regards to future TDP-43 investigations, our findings highlight the need to employ tissue models, which are most relevant for a certain condition. Most importantly, our results show that in the case of TDP-43 proteinopathies the knowledge acquired by studying neuronal cells could be translated to muscles only to a limited extent. Despite not investigated in this work, the same presumably applies for the interpretation of iCLIP results, in which TDP-43 binding should be always considered in the context of tissue characteristic environment, having in mind possible differences in binding behaviour of the protein across cell types that might affect splicing (Highley *et al*, 2014) and expression changes (Klim *et al*, 2019).

In some cases, our parallel study has given expected results. In NSC34 cells, for example, TDP-43 loss impacts expression of genes participating in pathways that provide elemental functions of neuronal cells, like *vesicle-mediated transport* and *regulation of postsynaptic membrane neurotransmitters*, which is perfectly in line with previous studies (Polymenidou *et al*, 2011; Tollervey *et al*, 2011). Similarly, the loss of TDP-43 in C2C12 cells impairs muscle characteristic features, like *striated muscle development, muscle cell migration* or *regulation of muscle cell differentiation*, what has been functionally confirmed by others(Militello *et al*,2018; Vogler *et al*, 2018). However, we have also detected tissue-specific TDP-43-associated dysregulation of molecular functions that will probably deserve further investigation. For example, we found that in muscles TDP-43 mediates splicing of mRNAs encoding proteins implicated in DNA-related processes. This is a particularly interesting observation as DNA-related processes play an important role in muscle differentiation. In adult skeletal muscle, DNA and histone modifications participate in adaptive response to environmental stimuli, which challenge structural and metabolic demands and thus make skeletal muscle a very plastic tissue (Barrès *et al*, 2012; McGee & Hargreaves, 2011). Also the early commitment towards myogenic lineage involves epigenetic changes mediated by chromatin remodelling enzymes like histone deacetylases (HDACs), histone acetyltransferases (HATs) and histone methyltransferases (HMTs) (Guasconi & Puri, 2009). In keeping with this, *Dnmt3a, Dnmt3b, Hdac9, Hdac7, Prdm2* are just few of chromatin-modifying enzymes that underwent splicing changes upon TDP-43 depletion in C2C12 but not in NSC34 cells. Interestingly, telomere shortening was described in primary muscle cultures of sIBM patients suggesting premature senescence (Morosetti *et al*, 2010) and epigenetic changes have been described in congenital myopathies (Rokach *et al*, 2015). Therefore, the results obtained in C2C12 suggest another possible mechanism on how TDP-43 may control gene expression in muscle in an indirect fashion and eventually participate in disease. Recently, loss of TDP-43 was associated with increased genomic instability and R-loop formation (Giannini *et al*, 2020; Wood *et al*, 2020) possibly through mechanisms involving Poldip3, which has been shown to play a role in maintaining genome stability and preventing R-loop accumulation at sites of active replication (Björkman *et al*, 2020).

On the other hand, some molecular processes such as dysregulation of protein assembly and disposal; mitochondrial changes and apoptosis; as well as alterations in post-translational modifications seem to occur upon TDP-43 depletion in both cell types, which possibly links these pathological changes to TDP-43 dysfunction in both tissues. As we have drawn our conclusions based on the RNA-seq analysis, a crucial future step will be to functionally assess to what extent TDP-43 loss impacts the above-mentioned processes in each tissue. Ideally, functional experiments should be performed in the two cell types in parallel, as only such approach would allow a direct comparison of the regulatory role played by TDP-43 in each context and would answer the question, whether impairment of RNA processing is as central in IBM as it is in ALS.

Working with cell lines representing muscles (Militello *et al*, 2018; Vogler *et al*, 2018; Reber *et al*, 2016) and neurons (Colombrita *et al*, 2009; Fiesel *et al*, 2012; Highley *et al*, 2014; Nonaka *et al*, 2009; Tollervey *et al*, 2011) allowed us a direct (and unbiased) assessment of TDP-43 activity across cell types. Mouse cell lines have been routinely employed to study TDP-43 (Militello et al., 2018; Vogler et al., 2018). In our case, they were chosen over human cells due to the lack of an appropriate and well-established cell line derived from human skeletal muscle. With regards to the contribution of TDP-43 malfunction to human pathology, we observed that transcripts, whose splicing was commonly affected by TDP-43 loss in the two mouse cell lines, appear more likely to undergo TDP-43-regulated processing also in human cells. Herein, we show for the first time that alternative sequences regulated by TDP-43 in both cell lines are better conserved between species than those regulated in a cell type-specific manner. Nonetheless, a conservation of the alternative sequence itself cannot guarantee for splicing conservation. Thus, it would be extremely insightful to investigate conservation of TDP-43-regulated splicing between human and mice on a transcriptome-wide level by actual sequencing experiment (rather than comparing gene sequences as such), possibly using analogous tissues (Cardoso-Moreira *et al*, 2020). A good example of commonly regulated event is skipping of exon 3 within *POLDIP3*, the regulation of which is conserved between mouse in humans (Fiesel *et al*, 2012; Shiga *et al*, 2012; Polymenidou *et al*, 2011) and has made it the most consistently detected event across studies. In this study, however, we identified two novel targets, ASAP2 and *PPFIBP1*, and show that they indeed undergo TDP-43-dependent splicing in all (mouse and human) cell lines tested. These additional findings could be of interest to identify common endpoints of mouse and human disease models that could then be used to monitor the efficiency of eventual novel therapeutic approaches or to follow disease course/onset.

Finally, as a *proof-of-principle*, we show that splicing alterations of TDP-43-dependent transcripts does in fact take place in different tissues (i.e., skeletal muscle and certain brain regions) affected by TDP-43 pathology. While expression levels of a given transcript heavily vary between individuals and, in our experience, seem to be influenced by experimental procedure itself (how and when biopsies are taken), the relative abundance of characteristic isoforms appears to be a more reliable readout. Considering cell-type-specific activity of TDP-43, it is reasonable to deduce that splicing of other TDP-43-controlled transcripts would be affected in the skeletal muscle and in neurons. In conclusion, we show that splicing changes as such indeed represent a robust indication of pathological conditions both in the skeletal muscle of IBM patients and in the brain of individuals affected with FTLD.

## MATERIALS AND METHODS

### Cell culture

C2C12 immortalized mouse myoblasts (ECACC), SH-SY5Y human neuroblastoma (ECACC) and RH-30 human rhabdomyosarcoma (kindly donated by Marc-David Ruepp) were maintained in DMEM (Thermo Fisher Scientific), supplemented with 10% FBS (Thermo Fisher Scientific) and antibiotics/antimycotics (Sigma-Aldrich) under standard conditions. NSC34 motoneuron-like mouse hybrid cell line (available in house) was cultured in DMEM (Thermo Fisher Scientific) with 5%FBS (Sigma-Aldrich) and antibiotics/antimycotics (Sigma-Aldrich). All experiments were performed with cells of similar passage number (± 2). To silence TDP-43 in C2C12 and NSC34 cells, 40 nM of siTDP (mouse siTDP 5’-CGAUGAACCCAUUGAAAUA-3’, Sigma-Aldrich) or non-targeting siLUC (5’-UAAGGCUAUGAAGAGAUAC-3’, Sigma-Aldrich) were mixed with 54 μl of RNAiMAX (Invitrogen) following the manufacturer’s reverse transfection protocol and applied to cells 700 000 seeded in a 10 cm dish. 48 h later, transfected cells were collected for subsequent analysis. The same reagent was used to silence TDP-43 in human SH-SY5Y and RH-30 cells. 400 000 RH-30 were seeded in a 60 cm dish, reversely transfected (human siTDP 5’-GCAAAGCCAAGAUGAGCCU-3’, Sigma-Aldrich or siLUC) and harvested 48 h later. To deplete TDP-43 in SH-SY5Y cells, 1 000 000 cells were seeded in a 6 cm dish and reversely transfected. After 48 h, they were transfected again and harvested 48 h later.

### Western blotting

Whole-cell extracts were resuspended in PBS in the presence of protease inhibitor and sonicated. 15 μg of protein sample were separated on a 10% Bis-Tris gels (Invitrogen) and transferred to the nitrocellulose membrane (Invitrogen). The membrane was blocked in 4% milk-PBST and proteins were stained using the following antibodies: anti-TDP-43 (rabbit, Proteintech, 1:1000), anti-GAPDH (rabbit, Proteintech, 1:1000), anti-HSP70 (rat, EnzoLife Science, 1:1000), anti-tubulin (mouse, available inhouse, 1:10000) and HRP-conjugated secondary antibodies anti-rabbit (goat, Dako, 1:2000), anti-mouse (goat, Dako, 1:2000), anti-rat (rabbit, Dako, 1:2000). Image acquisition and result quantification were conducted using Alliance Q9 Advanced Chemiluminescence Imager (UviTech).

### RNA extraction, RT-PCR

Total RNA was isolated using standard phenol-chlorophorm extraction. Only undegraded (RIS > 8) RNA of high purity (A260/A230 and A260/A280 > 1.8) was taken for subsequent analysis. 500 ng of RNA was reversely transcribed using random primers (Eurofins) and Moloney murine leukaemia virus reverse transcriptase (M-MLV, Invitrogen) according to manufacturer’s instructions.

### Splicing-sensitive PCR and qPCR

For detection of alternatively spliced mRNAs, PCR primers were designed complementary to constitutive exonic regions flanking the predicted alternatively spliced cassette exon. PCR mix was prepared using gene-specific primers (0.6 μM, Sigma, primer sequences in the **Appendix Table S1** and **S2**) and TAQ DNA polymerase (Biolabs or Roche) according to manufacturer’s instructions and subjected to 35-45 cycles long thermal protocol optimized for each primer pair. PCR products were separated by capillary electrophoresis (DNA screening cartridge, Qiaxcel) and splicing transitions were quantified using Qiaxcel software (QIAxcel ScreenGel (v1.4.0)). Exon inclusion was calculated by the software. Percentage of the inclusion (Inc. %) reports the area under the curve of the peak representing the longer (inclusion) splicing isoform. For assessment of transcript levels, real-time quantitative PCR was performed using PowerUp SYBR Green master mix (Applied Biosystems) and gene-specific primers (primer sequences in the **Appendix Table S3**). cDNA was subjected to 45 cycles of the following thermal protocol: 95 °C for 3 min, 95 °C for 10 s, 65 °C for 30 s, 95 °C for 10 s, 65 °C for 1 s. Relative gene expression levels were determined using QuantStudio design and analysis software (Thermofisher Scientific (v1.5.1)) always comparing treated samples (siTDP) with their direct controls (siLUC) normalized against *Gapdh*. p-values were calculated using one-tailed paired t-test as qPCR was conducted to validate expression changes detected by RNA-seq.

### RNA-seq

Both polyA cDNA library generation and RNA-seq were performed by Novogene (Beijing, China). cDNA libraries with insert length of 250-300 bp were generated using NEB NextUltra RNA Library Prep Kit. Sequencing was conducted on Illumina with paired-end 150 bp (PE 150) strategy.

### Read mapping

Sequencing quality control and filtering were performed to prune reads with average Phred score (Qscore) below 20 across 50% of bases, as well as those with more than 0.1% of undetermined (N) ones. Obtained reads were aligned to the mouse genome GRCm38 (mm10) using the Spliced Transcripts Alignment to a Reference (STAR) software (v2.5) (Dobin *et al*,2013), an RNA-seq data aligner that utilizes Maximal Mappable Prefix (MMP) strategy to account for the exon junction problem.

### Quantification of gene expression level

Counting of reads mapped to each gene was performed using HTSeq (v0.6.1) (Anders *et al*,2015). Raw read counts together with respective gene length were used to calculate Fragments Per Kilobase of transcript sequence per Millions base pairs sequenced (FPKM). In contrast to read counts, FPKMs account for sequencing depth and gene length on counting of fragments (Mortazavi *et al*, 2008) and are frequently used to estimate gene expression levels.

### Differential expression analysis

Differential gene expression (DEG) analysis of two conditions was performed using the *DESeq2* R package (v2_1.6.3) (Anders & Huber, 2010), a tool that utilizes negative binomial distribution model to account for variance-mean dependence in count data and tests for differential expression (Love *et al*, 2014). Three biological replicates were included per cell type and condition, in control (siLUC) and TDP-43-silenced (siTDP) cells. Read count matrix was pre-filtered by removing rows with row sum below one. Multiple testing adjustments were performed using Benjamini and Hochberg’s approach to control for the false discovery rate (FDR). Transcripts with p_adj_ < 0.05 were considered as differentially expressed.

Differentially expressed genes identified in both cell lines under different experimental conditions were hierarchically clustered based on log_10_(FPKM+1) and visualized with *pheatmap* R package (v1.0.12) (Kolde, 2019). Further, distance between silenced and control samples of each cell line was illustrated with principal component analysis (PCA), using the R function *“prcomp”* (R Core Team, 2019). Differences in gene expression levels (log_10_(FPKM+1)) between cell lines were tested for significance using Wilcoxon signed-rank test.

### Alternative splicing analysis

Five major types of alternative splicing events – skipped exons (SE), mutually exclusive exons (MXE), alternative 5’ and 3’ splice sites (A5’SS and A3’SS) and intron retention (RI) - were detected and analysed by Novogene using replicate multivariate analysis of transcript splicing (rMATS) software (v3.2.1) (Shen *et al*, 2014). Every alternative splicing event can produce exactly two isoforms. Each isoform is adjusted for its effective length before calculating the ratio of two isoforms and testing significance of differential splicing between two conditions. Multiple testing was corrected using Benjamini and Hochberg’s method. Splicing events having FDR < 0.01 were considered significant irrespective of ΔPSI.

Alternatively (for analysis of cryptic splicing and patient’s data (**Fig 3D** and **Fig 7** and **Appendix fig S4D**)), differential splicing analysis was performed using MAJIQ (v2.1) and the GRCm38 as a reference genome as previously described elsewhere (Brown *et al*, 2021).

### Enrichment analysis

Gene Ontology GO (Ashburner *et al*, 2000) and Kyoto Encyclopaedia of Genes and Genomes databases KEGG (Kanehisa, 2000) are widely used in gene enrichment analysis to classify list of individual genes based on their expression pattern, or other similar feature, with the aim to predict dysregulated biological processes, functions and pathways or any other general trend within a subset of data (Yu *et al*, 2012). In this study, GO enrichment and KEGG analysis were conducted in R, using *clusterProfiler* package (v3.14.3) (Yu *et al*, 2012) either on the set of differentially expressed genes (p_adj_ < 0.05) or alternatively spliced genes (FDR < 0.01), if not stated otherwise. Additionally, GO enrichment analysis was conducted using less stringent threshold for inclusion of alternatively spliced genes (where we considered genes with non-corrected p-value < 0.01 instead of FDR< 0.01). Genes of a particular dataset were assigned Entrez gene identifiers from Bioconductor mouse annotation package *org.Mm.eg.db* (v3.10.0). Enrichment test for GO terms and KEGG pathways were calculated based on hypergeometric distribution. The resulting GO terms/KEGG pathways were considered significant after applying multiple testing corrections with Benjamini-Hochberg method (p_adj_ < 0.05). Subsequently, significant GO terms (category: biological process) were functionally grouped or manually edited depending on the underlying biological question.

### Conservation analysis

Gene/exon conservation analysis within mouse (mm10) was performed by calculating phyloP (phylogenetic p-values) scores, i.e., per base conservation scores, generated from aligned genomic sequences of multiple species (Pollard *et al*, 2010).

For each differentially expressed gene, the average *per gene* phyloP score was computed with bigWigSummary (UCSC) (Kent *et al*, 2010). To calculate phyloP scores of TDP-43-regulated alternative sequences (hereafter referred to as *per exon* phyloP score), we considered TDP-43-regulated sequences of all event types. Those include A’3SS and A’5SS (long and short exon), retained introns and cassette exons (SE, the 1^st^ and the 2^nd^ exon of MXE).

### Patient samples

The NYGC ALS cohort has previously been detailed elsewhere (Prudencio *et al*, 2020; Brown *et al*, 2021). Herein, we only considered ALS and FTLD patients with TDP-43 pathology (ALS-TDP and FTLD-TDP) and healthy controls while excluding ALS with *SOD1* mutations of FTLD patients without TDP-43 inclusions.

Muscle biopsies (vastus lateralis or biceps) were obtained from 4 patient diagnosed with IBM according to the Griggs criteria (Griggs *et al*, 1995) and 4 healthy controls. Participants were investigated for cramps or fatigue, they underwent regular examination, neurophysiology tests and histological examinations. IBM biopsies were taken from moderately affected muscles and routinely investigated for histological and immunohistochemistry features. In case muscle fibrosis was present, it did not compromise a definite pathologic diagnosis. Basic demographic features of all participants are summarised in **Appendix Table S4**. Biopsies were stored at 80 °C. Institutional board reviewed the study and ethical approval was obtained.

Sample processing, library preparation, and RNA-seq quality control have already been described elsewhere (Brown *et al*, 2021).

## DATA AVAILABILITY

Datasets generated for this study are deposited in NCBI’s Gene Expression Omnibus and are accessible through GEO Series accession number GSE171714. [The following secure token has been created to allow review of record GSE171714 while it remains in private status: ibkdwqwkxjuvbaj].

## ACKNOWLEDGEMENTS

We thank Marc-David Ruepp (King’s College London) for providing RH-30 cells and Robert Bakarić for his kind assistance with the conservation analysis.

We would also like to thank the Target ALS Human Postmortem Tissue Core (New York Genome Center for Genomics of Neurodegenerative Disease, Amyotrophic Lateral Sclerosis Association) for providing post-mortem brain samples, patients and their families who donated those samples. See supplemental acknowledgments (**Appendix Table S5)** for the complete list of the NYGC ALS Consortium members. IBM muscle biopsies were kindly provided by the Bank of muscle tissue, peripheral nerve, DNA and Cell Culture, a member of Telethon network of Genetic biobanks, at Fondazione IRCCS Ca’ Granda, Ospedale Maggiore Policlinico, Milano, Italy and from the Laboratory of Muscle Histopathology and Molecular Biology at IRCCS Policlinico San Donato, San Donato Milanese, Italy.

This research was supported by the AriSLA grant PathensTDP to EB and by the European Reference Network for Neuromuscular Diseases to MM and MR. Consortium activities were supported by the ALS Association (15-LGCA-234) and the Tow Foundation. GM was supported by Fondazione Malattie Miotoniche, Milan, Italy. Andrea Cortese would like to thank Medical Research Council (MR/T001712/1), Cariplo foundation, the Italian Ministry of Health (Ricerca Corrente 2018–2019), the Inherited Neuropathy Consortium and the Fondazione Regionale per la Ricerca Biomedica for the grant support.

## AUTHOR CONTRIBUTIONS

UŠ and YA conducted experiments; UŠ, NŠ, ALB and MR analysed the data; UŠ and EB designed the study; HP provided patient data collected by the NYGC ALS consortium (see the complete list of members in the supplemental acknowledgments **Appendix Table S5**); AC, CC, EB, RC, GB, MR, MM provided IBM patient samples; PF, MS and EB supervised the study; UŠ and EB wrote the manuscript with contributions of other authors.

---

**Appendix figure S1.**
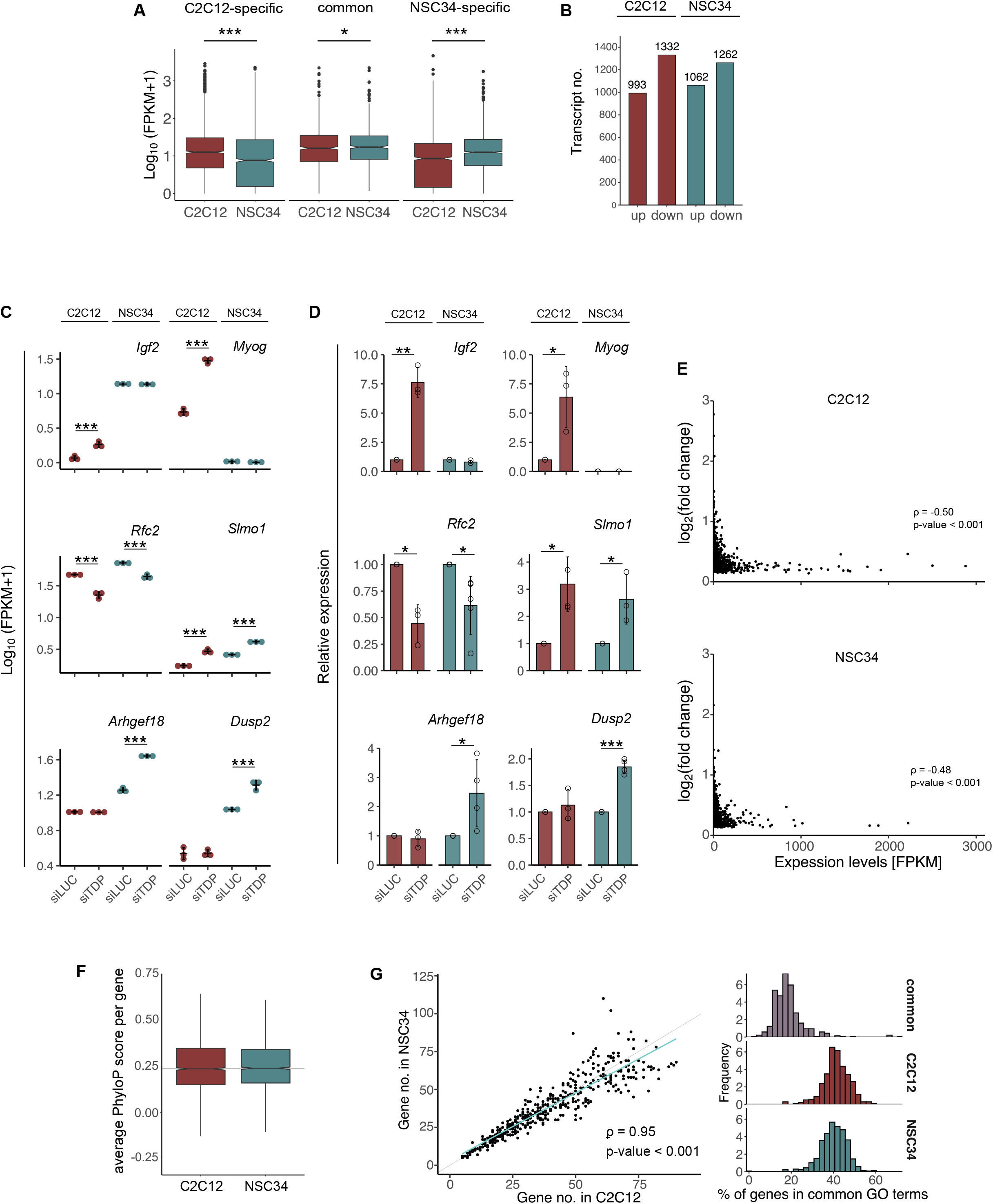
DEGs detected in C2C12 and NSC34. **A** The plot shows log_10_-transformed FPKM values of muscular, neuronal and common TDP-43 targets in siLUC-transfected C2C12 and NSC34 cells. C2C12-specific DEGs exhibit higher expression in C2C12 cells (p-value < 2.2·10^-16^), while NSC34-specific DEGs have higher expression in NSC34 (p-value < 2.2·10^-16^). Expression levels of common targets is more similar between cell lines (p-value = 0.02). Significance was tested using Wilcoxon signed-rank test. **B** The diagram shows the number of upregulated and downregulated genes detected in C2C12 and NSC34 cells following TDP-43 silencing. **C** Expression changes of representative DEGs (C2C12-specific vs. common vs. NSC34-specific, **Fig 2D**) as assessed by RNA-seq and plotted as log_10_-transformed FPKM. **D** Relative expression changes of DEGs from (**C**) were validated using qPCR. p-values were generated using Student’s t test (paired, one-tailed, n ≥ 3 per group). **E** Scatter plots show there is no correlation between the absolute change in gene expression following TDP-43 depletion (plotted as log2-transformed fold change) and the baseline expression of a given transcript (FPKM in siLUC-transfected cells) for DEGs identified in C2C12 (2325) and NSC34 (2324) (Spearman’s ρ = −0.50, p-value < 2.2·10^-16^ and Spearman’s ρ < −0.48, p-value < 2.2·10^-16^, respectively). **F** Average *per gene* PhyloP conservation scores plotted as box plots show TDP-43-regulated DEGs detected in C2C12 (2325) and NSC34 (2324) are equally well conserved across species (p-value = 0.48). p-value was generated using Wilcoxon rank sum test, the grey line represents the median of average PhyloP scores of all exons in the mouse genome. **G** The number of DEGs found in commonly enriched GO terms (**Fig 2E**, 459) is similar between two cell lines (left). Grey line represents y = x and the blue line the fitted regression (Spearman’s ρ = 0.95, p-value < 2.2 ·10^-16^). Frequency plot shows that commonly regulated terms are highly enriched for cell-type-specific TDP-43-regulated DEGs (right).

**Appendix figure S2.**
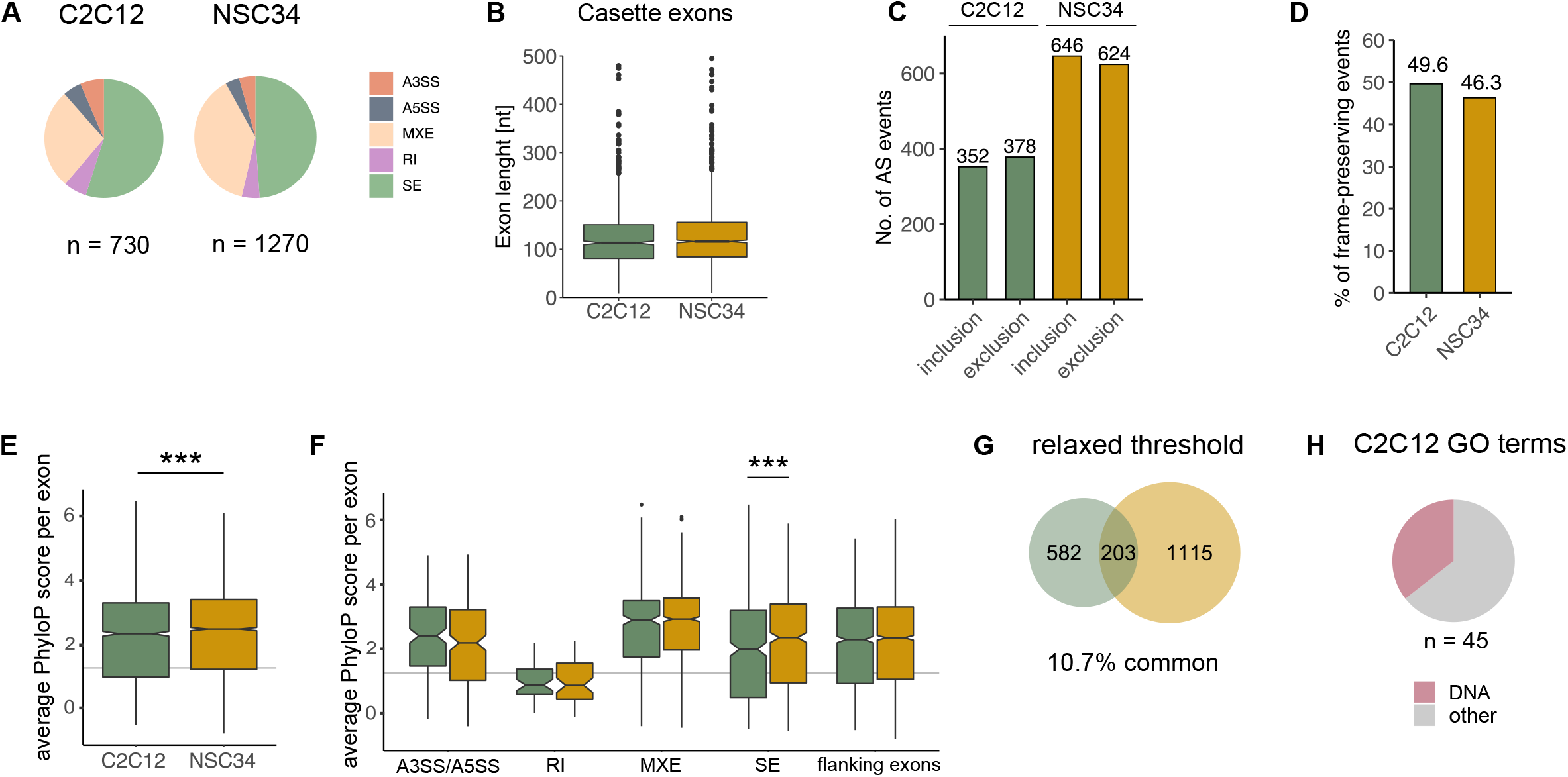
General features of TDP-43-controlled AS events detected in C2C12 and NSC34. **A** TDP-43-regulated AS events detected in C2C12 and NSC34 cells do not differ in terms of event type distribution (the number below shows the total number of AS events detected in each cell line); **B** the average length of TDP-43-regulated cassette exons (SE and MXE); **C** the ratio between inclusion/exclusion events; **D** the percentage of frame-conserving events. **E** Average *per exon* PhyloP conservation scores plotted as box plots show TDP-43-regulated alternative sequences detected in NSC34 cells (4281) are better conserved across species than those detected in C2C12 cells (2372) (p-value = 1.1·10^-4^). p-value was generated using Wilcoxon rank sum test with continuity correction, the grey line represents the median of average PhyloP scores of all exons in the mouse genome. **F** Average *per exon* PhyloP conservation scores of TDP-43-regulated alternative sequences stratified by event type (SE, MXE, RI, A3’SS, A5’SS). The difference (p-value = 6.5·10^-6^) among all groups was tested with Kruskal-Wallis rank sum test, followed by pairwise comparisons using Wilcoxon rank sum test with Benjamini-Hochberg correction for multiple testing. Significant difference is highlighted only for within event comparison between two tissues, SE (p-value = 9.7·10^-3^). The grey line represents the median of average PhyloP scores of all exons in the mouse genome. **G** Venn diagram shows the total number of AS events detected by rMATS at relaxed threshold (considering events that were detected at FDR < 0.01 in one dataset and p-value < 0.05 in the other). **H** GO enrichment analysis (refers to **Fig 3G**) was performed on alternatively spliced genes detected in C2C12 using less stringent threshold for genes which entered GO analysis (p-value < 0.01 instead of FDR < 0.01). Resulting GO terms (45) imply on dysregulation of DNA-related biological processes.

**Appendix figure S3.**
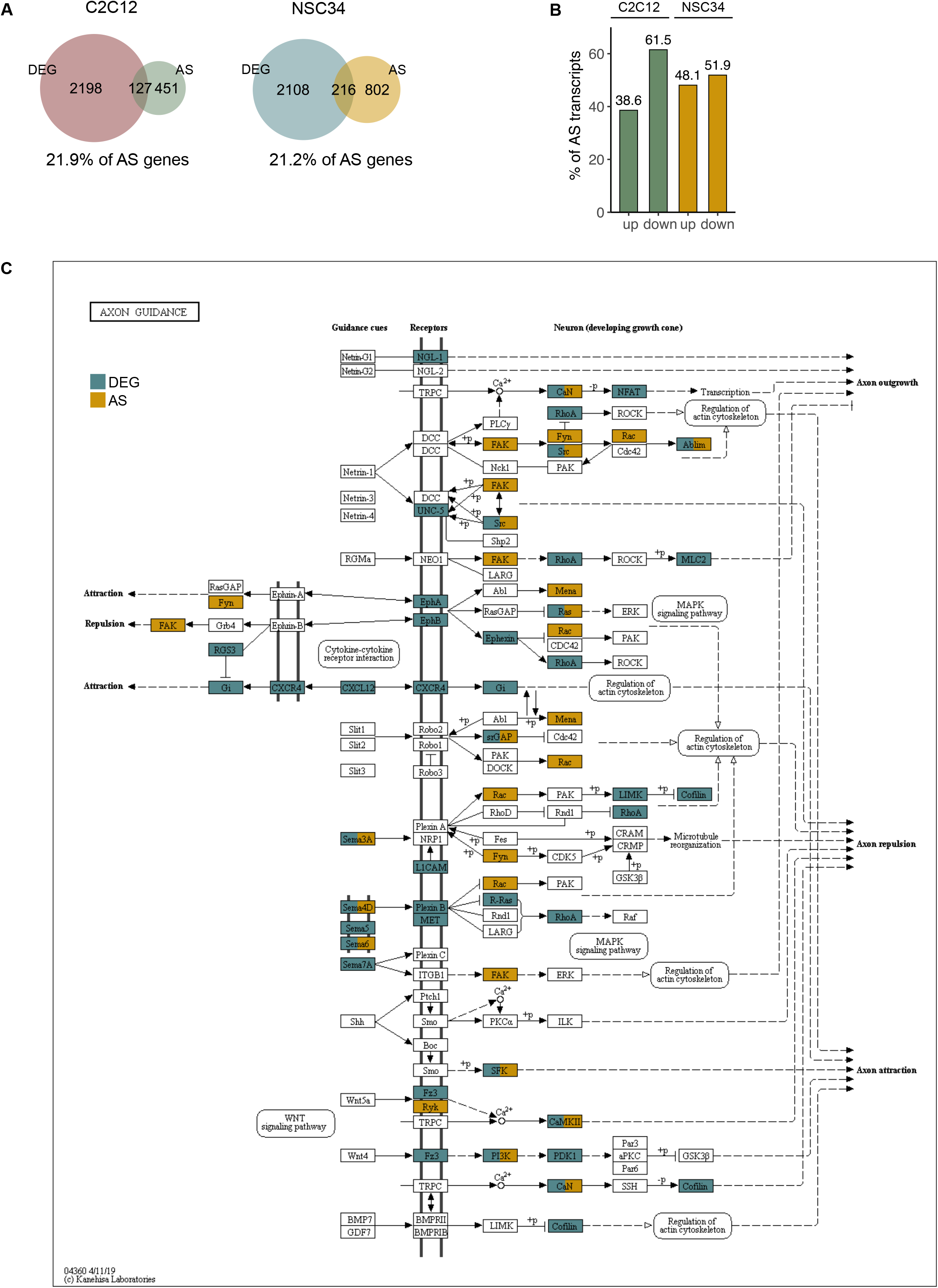
Transcripts subject to TDP-43-dependent splicing and expression level changes. **A** Venn diagrams show the percentage of transcripts affected by TDP-43 loss due to altered splicing (AS) or changes in the overall transcript abundance (DEG) in each cell line (21.9% of AS genes in C2C12 and 21.0% of AS genes in NSC34, respectively). **B** The barplot shows the percentage of down- and upregulated genes among those subject to altered splicing following TDP-43 loss. **C** A representative KEGG pathway (axon guidance pathway, mmu04360) significantly enriched (p_adj_ < 0.05) in AS and DE genes in NSC34 cells demonstrates that TDP-43 might influence axon guidance by regulating AS and expression levels of transcripts encoding proteins that participate in the given biological process. Proteins encoded by AS transcripts are shown in yellow and those encoded by DEG are shown in blue.

**Appendix figure S4.**
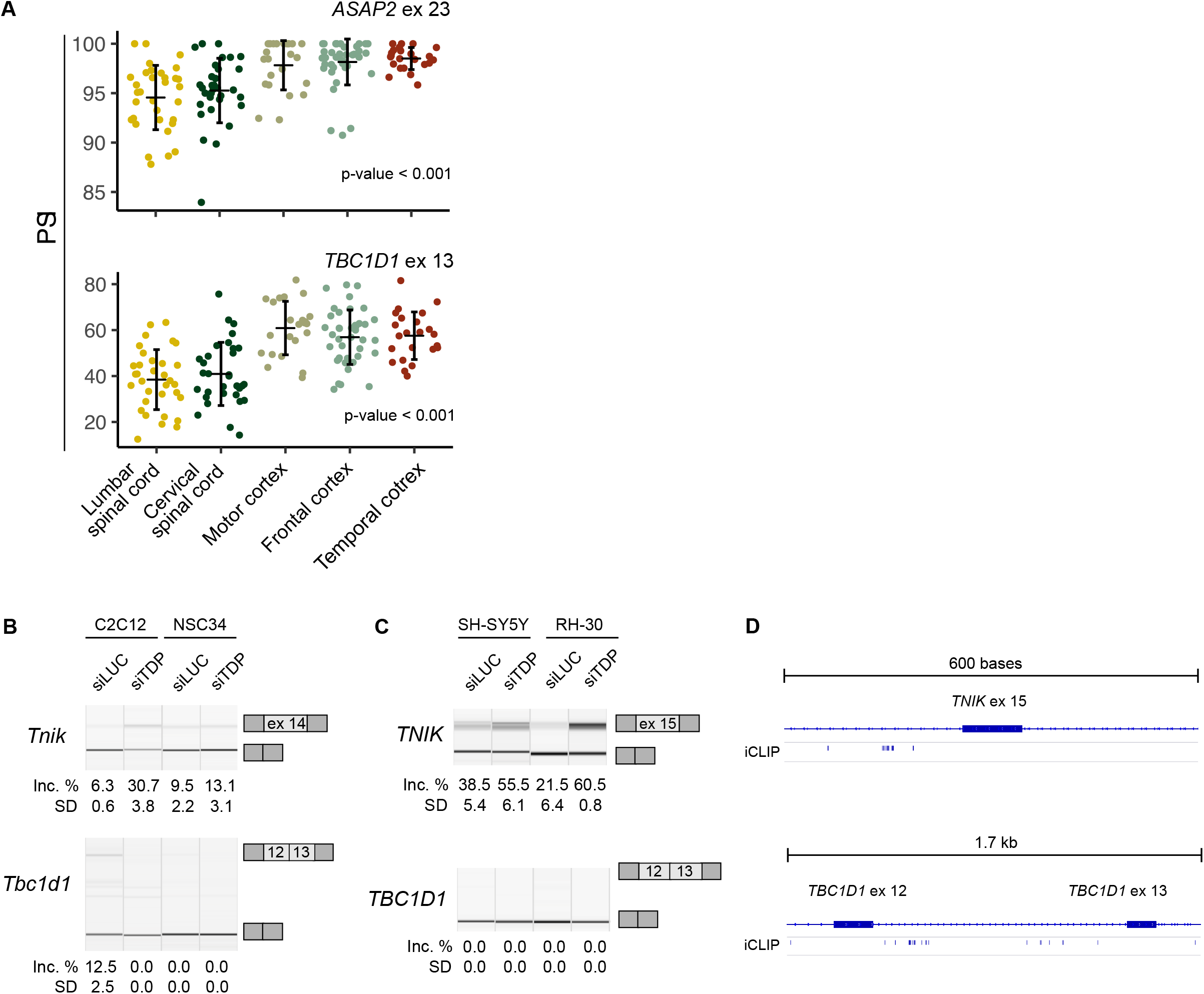
Tissue-characteristic inclusion of alternative exons regulated by TDP-43. **A** Dot plots demonstrate variable inclusion levels of alternative exons across different brain regions of healthy controls (in the absence of TDP-43 pathology), as exemplified by two alternative exons – exon 23 of *ASAP2* and exon 13 of *TBC1D1*. p-values (p-value = 1.6·10^-9^ for *ASAP2* and p-value = 2.4 ·10^-11^ for TBC1D1, respectively) were generated using Kruskal-Wallis chi-squared test. **B** TDP-43-depenent splicing of exon 14 of mouse *Tnik* and exons 12 and 13 of mouse *Tbcld1* occur in cell-type-specific fashion in mouse C2C12 and NSC34 cells. **C** Exon 15 of human *TNIK* is regulated by TDP-43 in both, SH-SY5Y and RH-30 cell line, likely in a direct fashion by TDP-43 binding in the upstream intron as shown in **(D)**. The long isoform of *TBC1D1* gene (exons 12 and 13 included) is not expressed in undifferentiated SH-SY5Y and RH-30 cells, as inclusion of exons 12 and 13 increases with differentiation (Bland *et al*, 2010), however, TDP-43 binding sites were identified in the vicinity of exons represented on panel **(D)**. (**B)**-**(C)** Semi quantitative RT-PCRs conducted in TDP-43-silenced cells and corresponding controls are shown along with the quantification of splicing changes (% of alternative exon inclusion) (see the exact transcript numbers in **Appendix Table S1** and **S2**, n = 3 replicates per group). **D** Schematic representation of TDP-43 binding sites identified by iCLIP analysis in SH-SY5Y cells (Tollervey *et al*, 2011) in the vicinity of exons represented on panel **(C)**.

